# A novel vimentin variant from tumor-associated macrophages directs cancer metastasis by engaging IGF-1R

**DOI:** 10.1101/2025.02.27.640678

**Authors:** Chenglin Miao, Xuedi Gao, Jingjing Mao, Wentong Wang, Zhifang Li, Zhiqin Chen, Zeyu Wen, Yuhan Huang, Xinyi Huang, Xin Liang, Xin Liu, Gaofeng Fan, Chenxi Tian, Suwen Zhao, Chengyuan Wang, Tao Xu, Yaming Jiu

## Abstract

Tumor-associated macrophages (TAMs) are recognized for their role in promoting malignancy. However, the specific mechanisms by which they drive cancer cell migration remain largely unexplored. Here, we identified a novel N-terminal-less variant of vimentin, a cytoskeletal intermediate filament protein, that is selectively secreted by TAMs through a type I unconventional secretion pathway following caspase cleavage of its N-terminus. The macrophage-secreted short vimentin (mssVIM) enhances tumor migration, as demonstrated in breast cancer cell lines, patient-derived cancer cells, mammospheres, and *in vivo* mouse xenograft models. Mechanistically, mssVIM binds and activates the insulin-like growth factor 1 receptor (IGF-1R) on the cancer cell surface, by exposing binding sites that are otherwise masked in full-length vimentin. Unlike the canonical IGF-1-IGF-1R signaling pathway that promotes cell proliferation, mssVIM-activated IGF-1R triggers ribosomal S6 kinase (RSK) activation, which increases integrin αVβ6 expression, ultimately facilitating tumor cell migration. Analysis of tissue samples from a cohort of breast cancer patients showed that mssVIM levels positively correlate with tumor malignancy and lymph node metastasis, suggesting its potential as a prognostic biomarker.

## Introduction

Breast cancer is the most commonly diagnosed malignancy and the leading cause of cancer-related deaths in women worldwide^1^. Metastasis is among the most fatal manifestations of breast cancer, significantly complicating treatment and substantially increasing mortality risk^2^. Primary management strategies for breast cancer include surgery, hormone receptor therapy, chemotherapy, and immune checkpoint therapy^3^. However, these treatments often face challenges due to limited effectiveness, frequent recurrence, resistance, and toxicity^4–6^, emphasizing the urgent need to uncover novel therapeutic targets.

Tumor microenvironment (TME) provides a nurturing niche for tumor progression^7^. Active extracellular factors within the TME contribute to tumor progression by degrading the extracellular matrix, promoting tumor cell proliferation, facilitating angiogenesis, and enhancing resistance to therapy^8–10^. Tumor-associated macrophages (TAMs) are strongly associated with increased metastasis, higher histological malignancy, and poor prognosis, making them a compelling therapeutic target^11–13^. Given that TAMs not only serve as key immune components but also directly drive tumor progression by secreting pro-tumorigenic factors^14–16^, identifying TAM-specific secreted oncogenic factors is crucial for understanding their role in cancer progression and for developing strategies to overcome therapeutic resistance.

As the most abundant type III cytoskeletal intermediate filament, vimentin is renowned for its well-established role in maintaining the structural integrity and mechanical resilience of cells^17,18^. It is also involved in cell migration, cell signaling, organelle positioning, and responses to cellular stress and wound healing^19,20^. While traditionally viewed as a cytoplasmic protein, vimentin is also found in extracellular spaces, such as the plasma and synovial fluid, where it contributes to inflammatory responses, pathogen defense, and angiogenesis, verified by the treatment of full-length recombinant vimentin protein^21–24^. Although secreted vimentin has been identified from monocyte-derived macrophages, endothelial cells and astrocytes^22,25,26^, its molecular characterization and function in cancer progression has remain elusive.

In this study, we demonstrated a novel paracrine communication from TAMs to breast cancer cells mediated by a unique form of cytoskeletal vimentin, termed mssVIM (macrophage-secreted short vimentin). This variant lacks the head domain due to caspase-3 cleavage and is selectively secreted via the type I unconventional secretion pathway. We uncovered that mssVIM interacts with the insulin-like growth factor 1 receptor (IGF-1R) on the surface of breast cancer cells, leading to the activation of downstream RSK-integrin signaling, ultimately promoting tumor metastasis both *in vitro* and in *vivo*. Notably, analysis of breast cancer tissue samples from patients revealed that mssVIM levels positively correlate with tumor malignancy and lymph node metastasis. Our findings shed light on the role of vimentin in accelerating cancer metastasis and its potential as a target for breast cancer treatment.

## Results

### Tumor-associated macrophages secrete N-terminal-less vimentin

Tumor interstitial fluid (TIF) is a critical representation of the complex TME landscape, encapsulating various secreted factors within the tumor milieu^27^. Mass spectrometry analysis of TIF from breast tumor tissues of human patients revealed cytoskeletal vimentin as the seventh most abundant protein (Fig. 1a, b, and Extended Data Fig. 1a). Comparison between tumor and peritumoral tissues revealed a significant increase in vimentin levels in TIF (Fig. 1b), which was further validated by Western blot analysis (Fig. 1c), potentiating the unreported functions of TIF vimentin in tumor progression. To investigate the source of vimentin, breast invasive carcinoma (BRCA) cells were extracted from patients’ tumor tissues, where vimentin was not detected in the supernatant (Fig. 1d, left panel). Detection of the supernatants from human cervical cancer HeLa and osteosarcoma U2OS cells further confirmed that vimentin was not derived from tumor cells (Extended Data Fig. 1b).

**Fig. 1.**
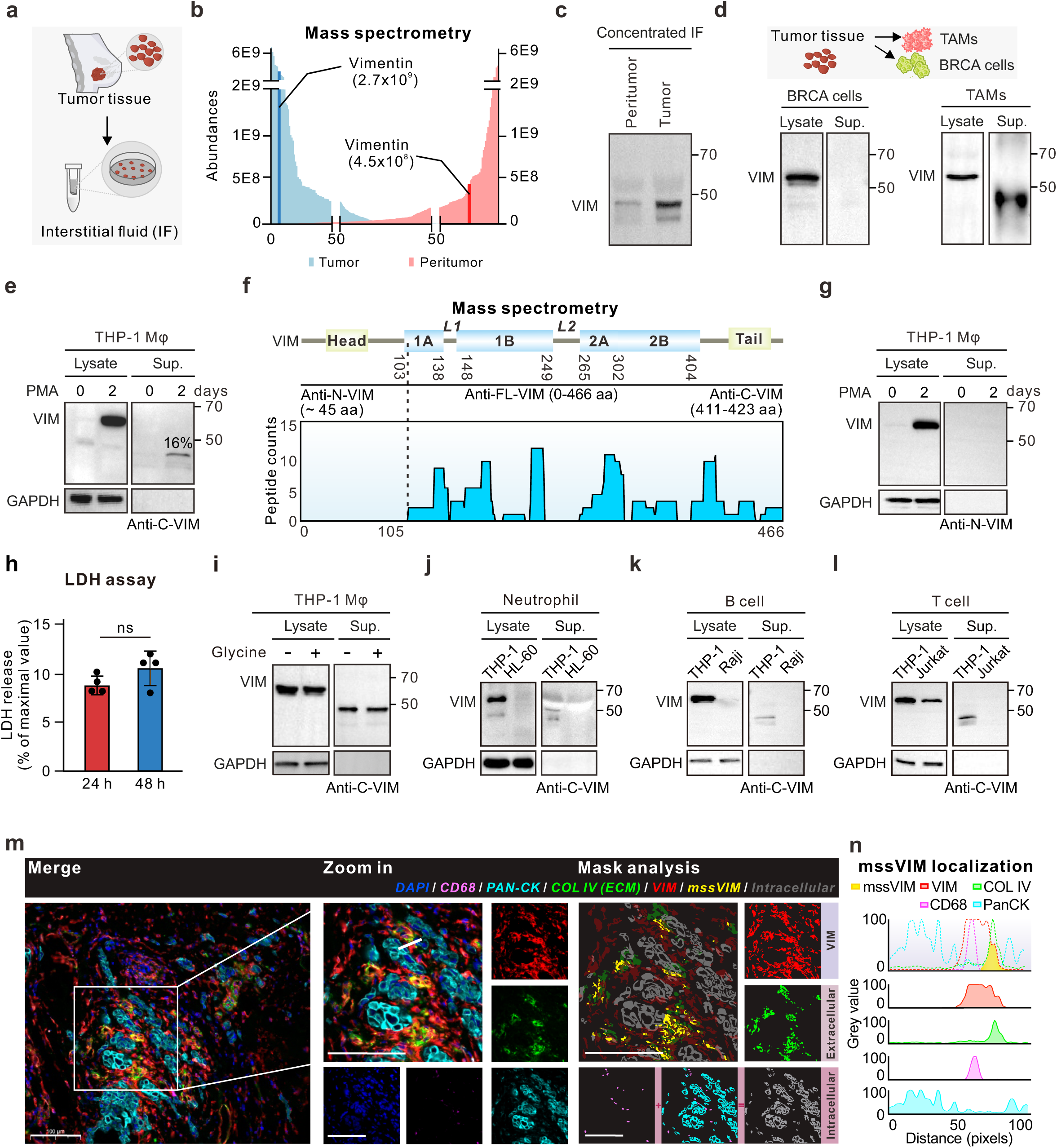
Tumor-associated macrophages (TAMs) exclusively secrete short-form vimentin. **a**, Schematic diagram depicting the interstitial fluid (IF) collection process from a breast invasive carcinoma (BRCA) patient. **b**, Quantification of the abundance of all top 500 proteins identified by mass spectrometry in the IF of tumor and peritumoral tissues. Blue columns represent tumors, while pink columns represent peritumoral tissue, the abundances of vimentin are 2.7×10^9^ and 4.5×10^8^, respectively. **c**, Western blot analysis of vimentin in the concentrated IF of the peritumor and tumor tissues from a BRCA patient. **d**, Schematic diagram depicting the processes of isolating tumor-associated macrophages (TAMs) and BRCA cells in patient tissues (upper panel). Western blot analysis of vimentin in the concentrated supernatants of BRCA and TAM cells (lower panels). **e**, Western blot analysis of vimentin immunoprecipitated from THP-1 differentiated macrophages (THP-1 Mφ) supernatant using an anti-C-terminal vimentin antibody (anti-C-VIM). GAPDH was used as a loading control. The ratio of secreted vimentin to cellular vimentin was 16%. **f**, Mass spectrometry of immunoprecipitated vimentin from the THP-1 Mφ supernatant showed the peptide counts of vimentin. The diagram in the upper panel shows the domain structure of vimentin and highlights the distinct recognition sites of three vimentin antibodies used in this study. **g**, Western blot analysis of vimentin immunoprecipitated from THP-1 Mφ supernatant using an anti-N-terminal vimentin antibody (anti-N-VIM). GAPDH was used as a loading control. **h**, The percentage of LDH released from THP-1 cells was measured at 24 h and 48 h using an LDH assay. **i**, Western blot analysis of vimentin in the cell lysate and the concentrated THP-1 Mφ supernatant treated with the plasma membrane rupture inhibitor glycine. GAPDH was used as a loading control. **j-l**, Western blot analysis of vimentin in the concentrated supernatants from HL-60 (**j**), Raji (**k**), and Jurkat (**l**) cells, respectively, with THP-1 Mφ supernatant as a positive control. GAPDH was used as a loading control. **m**, Representative fluorescent images of tumor tissues from a BRCA patient with antibody staining to visualize nucleus (DAPI), tumor cells (pan-CK), macrophages (CD68), extracellular matrix (collagen IV, Col IV), and vimentin (VIM). Mask analysis shows the distribution of macrophage-secreted short-vimentin (mssVIM) in yellow in the tumor microenvironment. Scale bars, 100 μm (left panel) and 50 μm (magnified images). **n**, Line profiles indicate the intensity of immunohistochemical staining based on the white line in Data are presented as mean ± S.D. from three independent experiments. ns P > 0.05; ***P < 0.001, ****P < 0.0001: Unpaired two-tailed Student’s t-test (**h**).

Analysis of the immune cells composition in metastatic breast cancer patients, based on the Cancer Genome Atlas Breast Invasive Carcinoma database (TCGA-BRCA), showed that tumor-associated macrophages (TAMs) are the most predominant immune cells in the TME (Extended Data Fig. 1c). Given the significant role of macrophages in tumor development and their association with decreased survival in breast cancer^14,28–30^, TAMs were isolated from breast cancer patients’ tissues. Remarkably, vimentin was detected in the supernatant (Fig. 1d, right panel). For further verification, we induced differentiation of the human monocyte cell line THP-1 into macrophages (Mφ), vimentin was also detected in the supernatant, which accounted for approximately 16% of the intracellular vimentin (Fig. 1e). Notably, instead of the typical 57 kDa full-length vimentin band (466 amino acids), distinct smaller bands at approximately 43 kDa were observed (Fig 1e). Similar results were observed in U937 Mφ cells (Extended Data Fig.1d). Mass spectrometry of the smaller vimentin bands from the supernatant of THP-1 Mφ revealed that the initial ∼105 amino acids (aa) at the N-terminus of vimentin were not detected (Fig. 1f), indicating that the vimentin in the Mφ supernatant may lack its N-terminal region. By employing antibodies targeting different epitopes of vimentin, we confirmed that vimentin in the supernatant could only be detected using the anti-C-terminal vimentin (anti-C-VIM) antibody (Fig. 1e). No signal was detected with the anti-N-terminal vimentin (anti-N-VIM) antibody (Fig. 1g).

To ascertain whether vimentin in the supernatant is derived from cell death, we measured the lactate dehydrogenase (LDH) release from THP-1 Mφ, as LDH is released into the culture medium during cell death. The results showed no significant increase in LDH release from THP-1 Mφ cells over a two-day period (Fig. 1h). Moreover, the application of the membrane rupture inhibitor glycine did not influence the vimentin levels in the supernatant of THP-1 Mφ cells (Fig. 1i)^31^. These findings imply that vimentin is actively secreted rather than passively leaked due to cell death.

To investigate whether other immune cells secrete vimentin, anti-C-VIM detection was applied to the supernatants from neutrophils, B cells, and T cells, which revealed no detectable bands (Fig. 1j-l). This result implied that vimentin in the supernatant is selectively secreted by macrophages. Therefore, we refer to this N-terminal-less vimentin as macrophage-secreted short vimentin, abbreviated as mssVIM.

To delineate mssVIM in the TME, we stained human breast cancer tissue samples with the extracellular matrix (ECM) marker collagen IV, tumor cell marker pan-CK, TAMs marker CD68, and anti-C-VIM, respectively. Although vimentin was visualized in the cytoplasm of both tumor cells and TAMs, it is worth noting the clear presence of extracellular vimentin surrounded by both tumor cells and TAMs (Fig. 1m). The intensity of immunohistochemical staining (represented as grayscale values) was evaluated to indicate the distribution of these markers across the tissues. Line profile analysis of fluorescence intensities for distinct markers further confirmed the presence of mssVIM in the TME (Fig. 1n).

### mssVIM is cleaved by caspase-3 and released via type I unconventional secretion pathway

To investigate whether mssVIM results from alternative splicing at the transcriptional level, we analyzed RNA sequencing data from wild-type (WT) THP-1 Mφ using Integrative Genomics Viewer (IGV). As a result, only full-length vimentin RNA was detected (Fig. 2a), inferring that mssVIM originates from cleavage at the protein level rather than alternative splicing. Given that calpain and caspases have been reported to cleave vimentin^32,33^, we assessed the impact of their inhibitors on the levels of mssVIM. The calpain inhibitor calpeptin had little impact on mssVIM levels (Extended Data Fig. 2a), whereas treatment with the pan-caspase inhibitor Z-VAD-FMK diminished mssVIM in the supernatant of THP-1 Mφ (Fig. 2b), suggesting caspases are involved in mssVIM cleavage. Consistent with reports that caspase-3 cleaves vimentin *in vitro* to produce a fragment of similar molecular weight^33^, the caspase-3 inhibitor Z-DEVD-FMK nearly abolished mssVIM production (Fig. 2c).

**Fig. 2.**
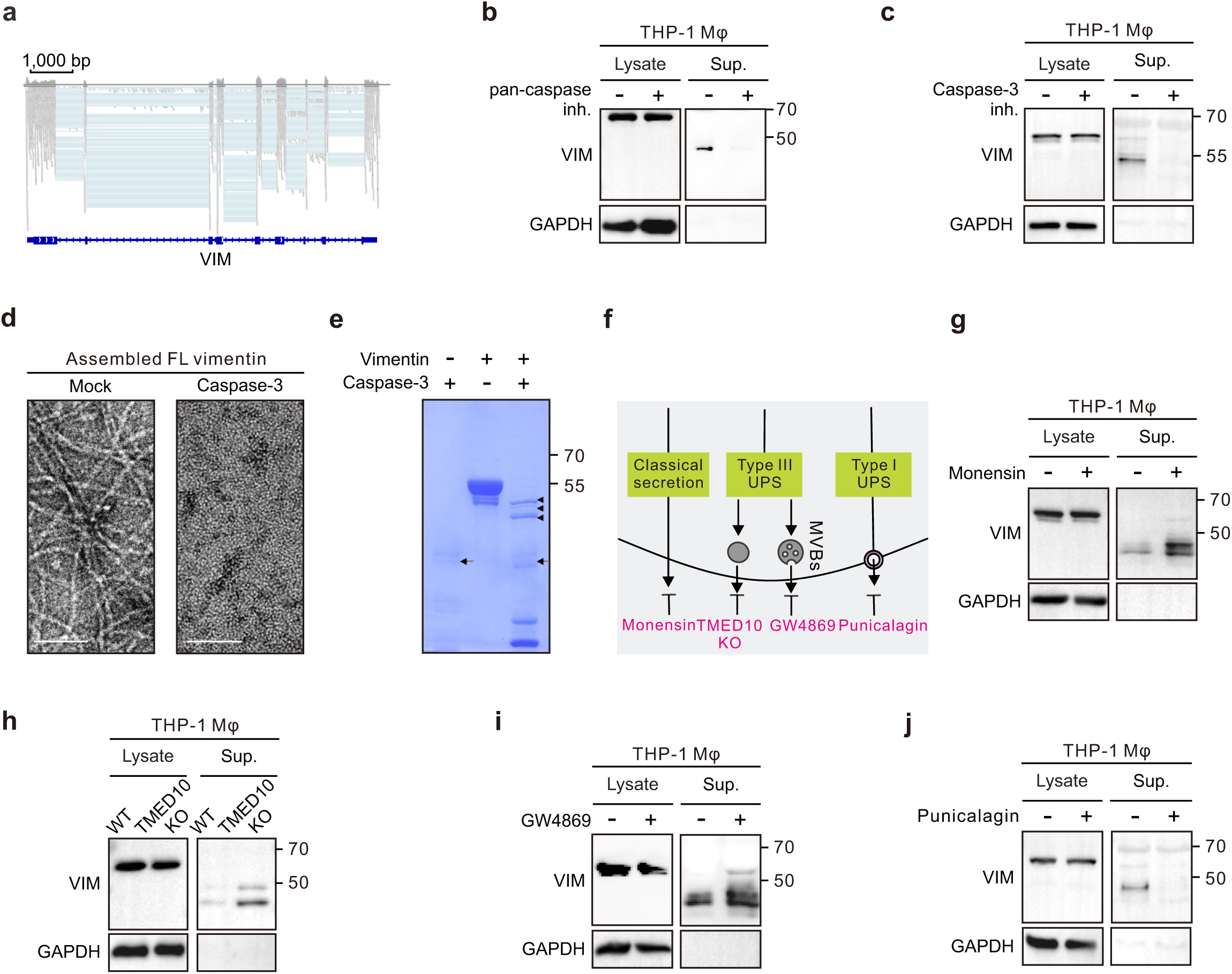
Vimentin is cleaved by caspase-3 and released via the type I unconventional protein secretion pathway. **a**, Integrative genomics viewer (IGV) analysis of THP-1 RNA sequence data indicates there is only one vimentin isoform at the RNA level. **b**, **c**, Western blot analysis of vimentin in the cell lysate and the concentrated THP-1 Mφ supernatants, treated with the pan-caspase inhibitor Z-VAD-FMK (**b**) or the caspase-3 inhibitor Z-DEVD-FMK (**c**). GAPDH was used as a loading control. **d**, Electron microscopy of negative staining images of recombinant assembled full-length vimentin filaments, with or without caspase-3 treatment. Scale bar, 100 μm. **e**, SDS-PAGE analysis of purified full-length vimentin incubated with or without caspase-3 protein. Black arrows indicate caspase-3 and black arrowheads indicate cleaved vimentin. **e**, Schematic illustration of recognized protein secretion pathways and corresponding specific inhibitors. **f**, Western blot analysis of vimentin immunoprecipitated from THP-1 Mφ supernatants, treated with or without monensin. **g**, Western blot analysis of vimentin immunoprecipitated from the supernatant of TMED10 KO THP-1 Mφ. **i**, **j**, Western blot analysis of vimentin immunoprecipitated from THP-1 Mφ supernatants, treated with or without GW4869 (**i**) or punicalagin (**j**).

To verify the cleavage of vimentin by caspase-3 in macrophages, electron microscopy was applied to visualize the *in vitro* filamentous network formation of recombinant vimentin. The network was disrupted by caspase-3 treatment, shown as sparse foci (Fig. 2d), indicating that head-cleaved vimentin no longer assembles into filaments. In a follow-up biochemical assay, recombinant vimentin was incubated with caspase-3 *in vitro*, resulting in three bands with molecular weights similar to mssVIM (Fig. 2e). Caspase-3 is known to cleave aspartic acid residues^34^. Given the molecular weight, multiple cleavage sites on vimentin are probable (Extended Data Fig. 2b). This is consistent with our results from the supernatants of TAMs and THP-1 Mφ, in which we also observed several smaller bands (Fig. 1c-e). These results confirm that mssVIM is derived from cytoplasmic filamentous vimentin and is cleaved by caspase-3.

To investigate how mssVIM is released from macrophages, we inhibited the known secretion pathways (Fig. 2f). Neither the inhibition of the classical secretion pathway by monensin nor the type III unconventional protein secretion (UPS) pathway by transmembrane Emp24 domain-containing protein 10 knockouts (TMED10 KO)^35^, nor the exosome secretion pathway by GW4869 prevent mssVIM secretion, as verified by blotting of vimentin from the supernatants of THP-1 Mφ (Fig. 2g-i). Instead, a compensatory increase in mssVIM was observed (Fig. 2g-i). Since vimentin lacks a signal peptide for extracellular secretion^22^, its release via the UPS is plausible. Type III UPS cannot influence mssVIM release, thus, we applied the type I UPS inhibitor punicalagin. As a result, punicalagin significantly reduced mssVIM levels in the supernatant of THP-1 Mφ (Fig. 2j). Together, these findings suggest that caspase-3 cleaves cytoplasmic vimentin inside macrophages to produce short vimentin, which is then secreted via the type I UPS.

### mssVIM triggers cancer cell migration *in vitro* and *in vivo*

To investigate the potential function of mssVIM in cancer progression, we established an *in vivo* breast cancer migration model using NOG female mice. Luciferase-expressing MCF-7 cells, a relatively low-invasive tumor cell line broadly used in breast cancer study^36^, were injected intravenously (I.V.), and specific treatments were applied to each group of mice. As a result, mice treated with THP-1 Mφ supernatant exhibited higher lung metastasis, indicated by increased luminescence intensity, compared to the control group and Anti-C-VIM co-treated group (Fig. 3a). Additionally, the lungs of mice treated with THP-1 Mφ supernatant showed an increased number of lesions and a more irregular surface, along with a higher proportion of metastatic areas observed by hematoxylin-eosin (H&E) staining (Fig. 3b). These effects were restored upon the addition of anti-C-VIM in the supernatant.

**Fig. 3.**
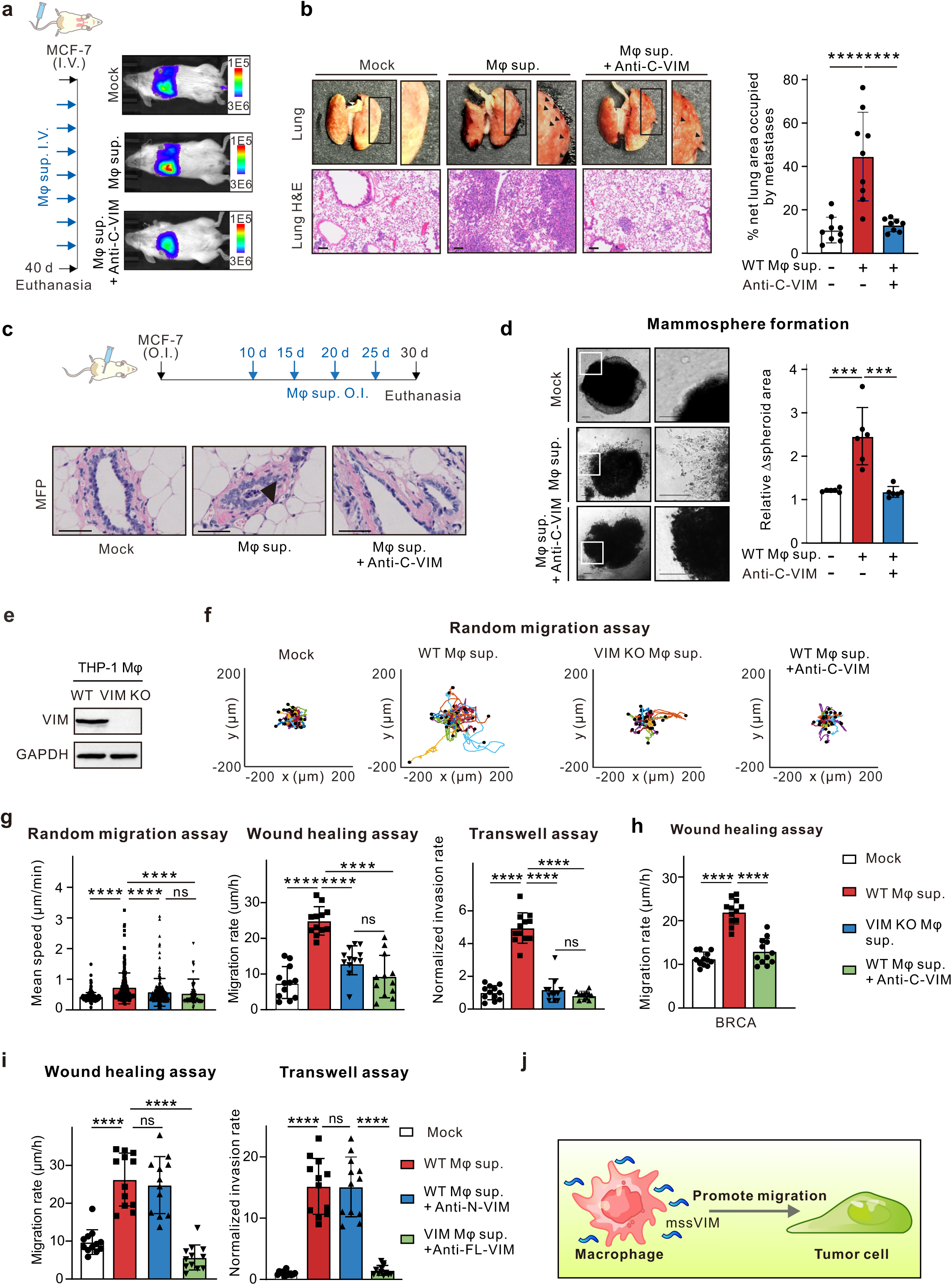
mssVIM promotes cancer cell migration *in vitro* and *in vivo*. **a**, Schematic diagram of the tail vein intravenous injection (I.V.) mouse model (left panel). Representative bioluminescence images of NOG mice showing luciferase metastasis to the lung (right panel). **b**, Representative images of whole mouse lungs and H&E staining of pulmonary nodules from mice intravenously injected with MCF-7 cells, treated with or without anti-C-VIM (left panel). Quantification of the percentage net lung area occupied by metastases (right panel). Scale bars, 100 μm **c**, Schematic diagram of the breast cancer mouse model with orthotopic injection (O.I.) of MCF-7- luciferase cells into nude mice (upper panel). H&E staining images show the level of local invasion in *in situ* tumors treated with the THP-1 Mφ supernatants from WT or WT with anti-C-VIM (lower panel). **d**, Representative images of MCF-7 mammosphere invasion process treated with THP-1 Mφ supernatants, with or without anti-C-VIM (left panel). Quantification of fold change in the mammosphere area (right panel). Scale bars, 20 μm. **e**, Western blot analysis of vimentin expression in wild-type (WT) and vimentin knockout (VIM KO) THP-1 Mφ. GAPDH was used as a loading control. **f**, Representative random migration trajectories of MCF-7 cells treated with WT or VIM KO THP-1 Mφ supernatants, with or without anti-C-VIM. **g**, Quantification of the mean speed in the random migration assay (left panel), migration rate in the wound healing assay (middle panel), and invasion rate in the transwell assay (right panel) of MCF-7 cells treated with WT or VIM KO THP-1 Mφ supernatants with or without anti-C-VIM. **h**, Quantification of the migration rate in the wound healing assay of BRCA cells treated with THP-1 Mφ supernatants with or without anti-C-VIM. **i**, Quantification of migration rate in the wound healing assay (left panel) and the invasion rate in the transwell assay (right panel) of MCF-7 cells treated with THP-1 Mφ supernatants, with or without anti-N-VIM or anti-FL-VIM. **j**, Schematic diagram of macrophages secreting mssVIM to promote tumor cell migration. Data are presented as mean ± S.D. from three independent experiments. ns P > 0.05; ***P < 0.001, ****P < 0.0001: One-way ANOVA Multiple Comparisons (**c**-**e**, **h**, **f**).

We further established an orthotopic injection (O.I.) breast cancer mouse model by injecting MCF-7 cells into the mammary fat pads of nude mice. THP-1 Mφ supernatants were administered every 5 days post-tumor cell inoculation. Immunohistochemistry on day 30 revealed that only the WT THP-1 Mφ supernatant induced breast cancer cell invasion (Fig. 3c), suggesting that mssVIM promotes cancer cell migration *in vivo*.

To quantify the invasive properties of MCF-7 cells, we developed a mammosphere invasion model. After a 5-day formation period and an additional 4 days for invasion within a matrigel-coated matrix, the WT THP-1 Mφ supernatant significantly increased the invasive levels, as shown by stellate morphology with spike-like invasive projections, which can be efficiently suppressed by combination with anti-C-VIM (Fig. 3d).

To elaborate the function of mssVIM, we generated a vimentin knockout (VIM KO) THP-1 cell line (Fig. 3e). No significant changes were observed in the concentration of total secreted proteins (Extended Data Fig. 3a). We conducted functional experiments, no significant difference in tumor proliferation was found when comparing the supernatants from wild-type (WT) and VIM KO THP-1 Mφ (Extended Data Fig. 3b), indicating that mssVIM did not affect tumor proliferation. Then we investigated whether mssVIM directly promotes cancer cell migration using wound healing, random migration, and transwell assays, which reflect the collective migration capacity, random migration ability, and invasion ability of tumor cells, respectively. As a result, the supernatant from WT THP-1 Mφ, but not from VIM KO cells, significantly increased the motility of MCF-7 cells, while addition of the anti-C-VIM antibody to the WT THP-1 Mφ supernatant inhibited its migration-promoting effect on MCF-7 cells in all three assays (Fig. 3f, g and Extended Data Fig. 3c, d). Similar effects were observed with the application of supernatant from U937 Mφ (Extended Data Fig. 3e). Importantly, the migratory capacity of BRCA cells isolated from patients’ tumor tissues was also enhanced by the application of WT THP-1 Mφ supernatant and reduced by the addition of anti-C-VIM antibody (Fig. 3h). This phenotype was further validated in the human cervical cancer cell line HeLa and the osteosarcoma cell line U2OS (Extended Data Fig. 3f). Further, dose-dependent neutralization experiments with anti-C-VIM further confirmed the function of mssVIM in promoting tumor cell migration (Extended Data Fig. 3g). Notably, using anti-full-length (FL)-VIM antibody, but not anti-N-VIM antibody, neutralized the pro-migration effect of the supernatant of WT THP-1 Mφ (Fig. 3i). Additionally, FL recombinant vimentin failed to replicate the pro-migration function of mssVIM (Extended Data Fig. 3h), implying the structural and functional differences between mssVIM and full-length vimentin. Together, our results suggest that mssVIM promotes cancer cell migration both *in vivo* and *in vitro* (Fig. 3j).

### mssVIM interacts with and activates IGF-1R in cancer cells

Cytoplasmic vimentin is a key marker of epithelial-mesenchymal transition (EMT) and contributes to enhancing cancer cell migration^37^. Given the low endogenous expression of vimentin in MCF-7 cells, we investigated the potential of mssVIM to enter the cytoplasm of MCF-7 cells and perform EMT function. Incubation with the supernatant from WT THP-1 Mφ did not induce visible vimentin signaling in MCF-7 cells. We then generated an MCF-7 cell line stably overexpressing vimentin (MCF-7 VIM OE) (Extended Data Fig. 4a, b). However, wound healing and transwell assays revealed that MCF-7 VIM OE cells did not exhibit increased migration and invasion abilities compared to WT MCF-7 cells (Extended Data Fig. 4c and 4d).

Subsequently, we hypothesized that mssVIM exert its pro-migration effects by activating membrane receptors. To identify the receptor activated by mssVIM, we performed biotinylation and enrichment of cell membrane surface proteins, followed by global phospho-proteomic mass spectrometry (Fig. 4a). The top abundant tumor-associated receptors were listed (Fig. 4b). We then predicted the binding affinity and dissociation constants of head-less vimentin (105-466 aa) with the receptors identified by mass spectrometry using AlphaFold (Fig. 4c and Extended Data Fig. 4e). The insulin-like growth factor 1 receptor (IGF-1R) was selected because it has a high binding affinity with head-less vimentin and it is significantly expressed in breast cancer cells. The activation level of IGF-1R was assessed with Western blot. Treatment with WT, but not VIM KO THP-1 Mφ supernatants significantly upregulated the phosphorylation level of IGF-1R, which can be neutralized by applying anti-C-VIM (Fig. 4d).

**Fig. 4.**
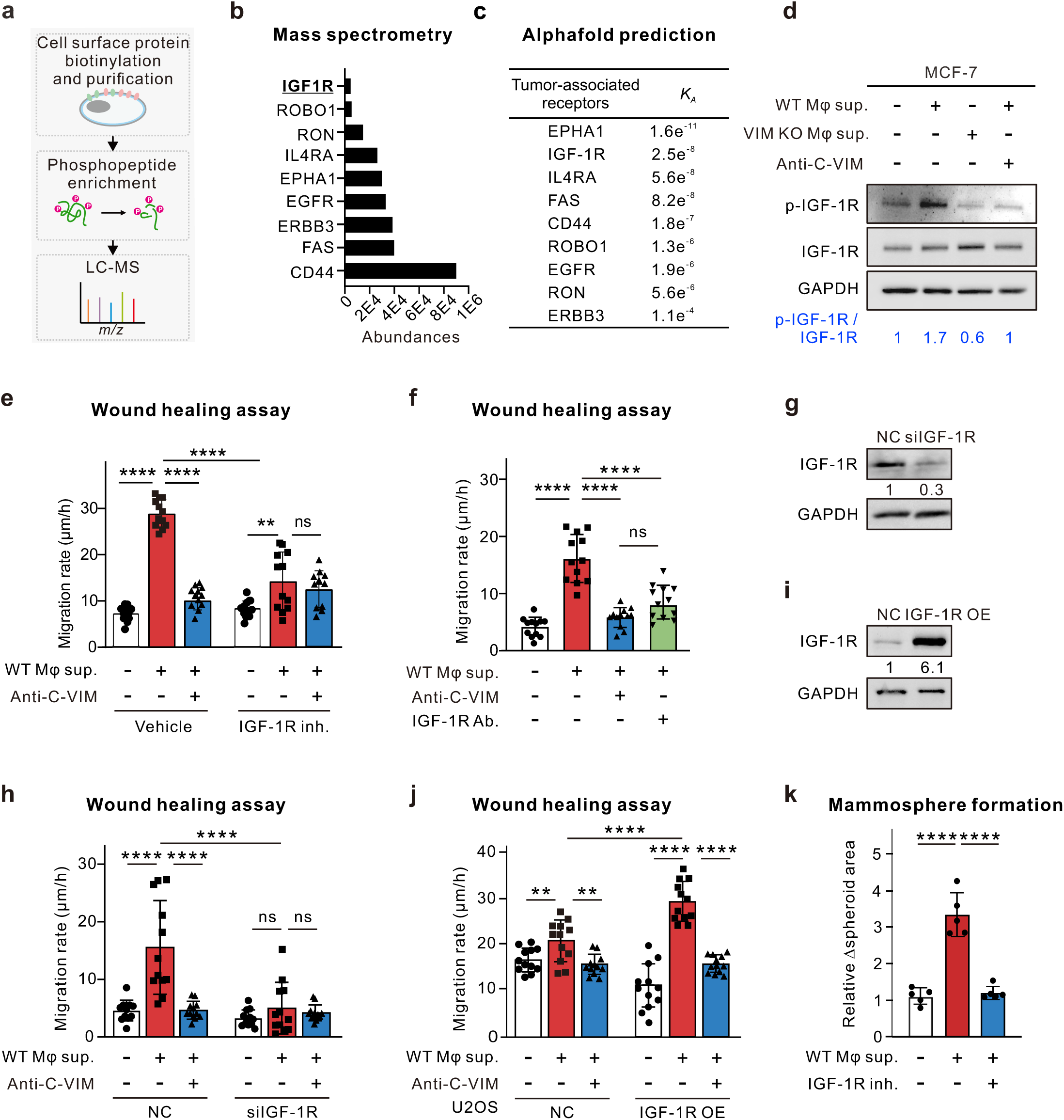
mssVIM interacts with and activates IGF-1R in tumor cells. **a**, Schematic diagram of the screening process for potential mssVIM targets by biotinylation-dependent cell surface protein extraction, phosphopeptide enrichment, and mass spectrometry. **b**, Quantification of the top abundant tumor-associated receptors identified by mass spectrometry depicted in (**a**). **c**, Binding affinity and dissociation constants of the heterodimeric structure of headless vimentin (105-466 aa) with the receptor from Fig. 4b, predicted using AlphaFold 3. **d**, Western blot analysis of p-IGF-1R and IGF-1R levels in MCF-7 cells treated with THP-1 Mφ supernatants, with or without anti-C-VIM. GAPDH serves as a loading control. The ratio of p-IGF-1R to IGF-1R is indicated by blue numbers. **e**, Quantification of migration rates in wound healing assay of MCF-7 cells treated with vehicle or IGF-1R inhibition, followed by treatment of THP-1 Mφ supernatants, with or without anti-C-VIM. **f**, Quantification of the migration rate in wound healing assay of MCF-7 cells treated with THP-1 Mφ supernatants, with or without anti-C-VIM or IGF-1R antibody. **g**, Western blot analysis of IGF-1R levels in MCF-7 cells, comparing the siRNA negative control (NC) and IGF-1R knockdown (siIGF-1R). GAPDH was used as a loading control. Numbers in the blots indicate the levels of IGF-1R expression normalized to GAPDH and the control group. **h**, Quantification of the migration rate in wound healing assay of NC and siIGF-1R MCF-7 cells treated with THP-1 Mφ supernatants, with or without anti-C-VIM. **i**, Western blot analysis of IGF-1R levels in U2OS cells, comparing the negative control (NC) and IGF-1R overexpression (IGF-1R OE). GAPDH was used as a loading control. Numbers in the blots indicate the levels of IGF-1R expression normalized to GAPDH and the control group. **j**, Quantification of the migration rate in wound healing assay of U2OS cells or U2OS IGF-1R OE cells treated with THP-1 Mφ supernatants, with or without anti-C-VIM. **k**, Quantification of fold changes in mammosphere area treated with THP-1 Mφ supernatants, with or without IGF-1R inhibitor PPP. Data are presented as mean ± S.D. from three independent experiments. ns *P* > 0.05; \*\*\**P* < 0.001, \*\*\*\**P* < 0.0001: One-way ANOVA multiple comparisons (**f**, **k**) and two-way ANOVA multiple comparisons (**e**, **h**, **j**).

Building on these findings, we conducted IGF-1R exogenous interventions in MCF-7 cells, including treatment with the IGF-1R inhibitor Picropodophyllin (PPP), neutralization with an IGF-1R antibody, and IGF-1R knockdown using small interfering RNA (siIGF-1R). All these treatments significantly inhibited the pro-migration effects induced by THP-1 Mφ supernatant (Fig. 4e-h, and Extended Data Fig. 4f, g). Notably, overexpression of IGF-1R in U2OS cells, which naturally exhibit low IGF-1R levels, obviously enhanced the pro-migration effect of mssVIM (Fig. 4i, j). Further, the IGF-1R inhibitor also inhibited the pro-migration function by THP-1 Mφ supernatant treatment in the mammosphere formation assay, which mimics *in vivo* condition (Fig. 4k). Together, these results confirm the critical role of IGF-1R in mssVIM-mediated promotion of cancer cell migration.

### The cleavage of the vimentin N-terminus facilitates the interaction between mssVIM and IGF-1R

The finding that the pro-cancer migration function of mssVIM is an IGF-1R-dependent process implies that mssVIM may bind to IGF-1R potentially like a ligand. We first used AlphaFold to predict the binding profile of head-less vimentin (105-466 aa) with the IGF-1R extracellular domain (ECD, 1-906 aa). The residues Tyr 319, Ser 325, Glu 338, and Ser 339 on head-less vimentin exhibited high affinity with the residues His 392, Asp 424, Gln 426, Thr 748, and Cys 544 on IGF-1R (Fig. 5a). These binding sites displayed a relatively high predicted local distance difference test (pLDDT) score (Extended Data Fig. 5a), indicating strong confidence and accurate predictions. Further, we performed a pull-down assay which verified the direct interaction between mssVIM in the THP-1 supernatant and His-tagged recombinant IGF-1R protein (Fig. 5b). To further verify the interaction between mssVIM and IGF-1R, we mutated the predicted five interacting sites in IGF-1R (IGF-1R Mut) and introduced the IGF-1R Mut in MCF-7 IGF-1R knockdown cells. As a result, the migration rate cannot be recovered by IGF-1R Mut compared to cells re-expressing WT IGF-1R (Fig. 5c).

**Fig. 5.**
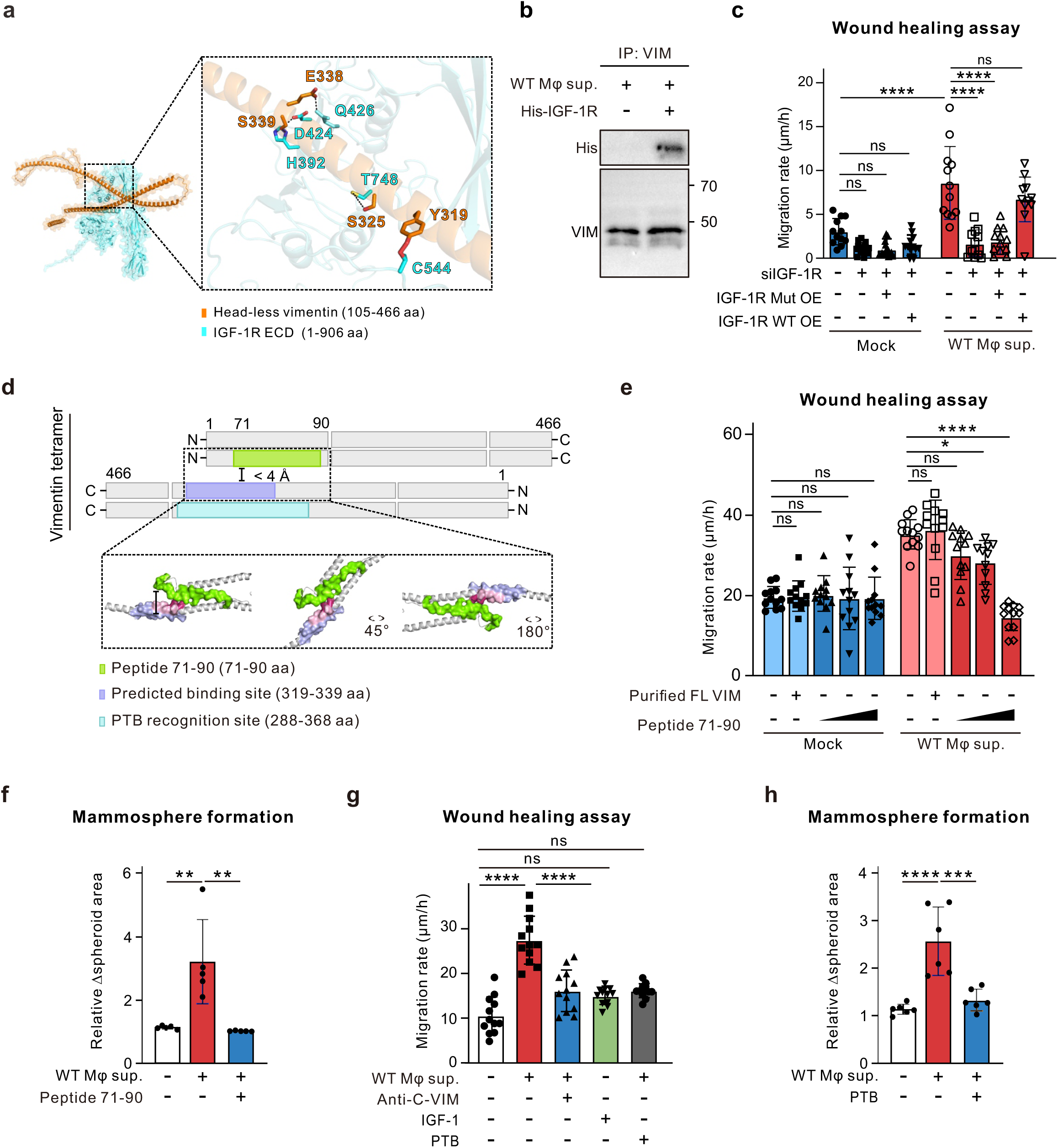
mssVIM binds to IGF-1R. **a**, AlphaFold predicts the interaction between head-less vimentin (blue) and the extracellular domain (ECD) of IGF-1R (orange). Dashed lines indicate hydrogen bonding or salt bridges between atoms <4.0 Å. Predicted residues participating in the binding are labeled, with vimentin colored blue and IGF-1R colored orange. **b**, *In vitro* Co-immunoprecipitation (Co-IP) assay followed by western blotting to test the association of recombinant His-tagged IGF-1R protein with mssVIM, which was enriched using anti-C-VIM from the concentrated THP-1 Mφ supernatant. **c**, Quantification of the migration rate of siNC and siIGF-1R MCF-7 cells in the wound healing assay, followed by overexpression of IGF-1R mutant or IGF-1R WT, treated with THP-1 Mφ supernatants, with or without anti-C-VIM. **d**, Diagram illustrating the presumed masking (distance <4.0 Å) within the vimentin staggered tetramer. Amino acids 71-90 and 319-339 in the vimentin head domain and rod domain are colored green and purple, respectively. The epitopes of the pritumumab (PTB) are colored blue. **e**, Quantification of the migration rate of MCF-7 cells treated with the supernatants from WT THP-1 Mφ, combined with purified FL VIM or increasing concentrations of vimentin peptide71-90 aa, in the wound healing assay. **f**, Quantification of fold changes in the mammosphere area treated with THP-1 Mφ supernatants, with or without vimentin peptide71-90 aa. **g**, Representative images of MCF-7 mammosphere formation treated with medium or THP-1 Mφ supernatants, with or without PTB. Scale bars, 20 μm. **h**, Quantification of fold changes in the mammosphere area treated with THP-1 Mφ supernatants, with or without PTB. Data are presented as mean ± S.D. from three independent experiments. ns P > 0.05; ***P < 0.001, ****P < 0.0001: Unpaired two-tailed Student’s t-test (**i**), one-way ANOVA Multiple Comparisons (**f**, **g**, **h**), two-way ANOVA Multiple Comparisons (**c**, **e**).

Furthermore, docking analysis of the full-length vimentin tetramer with IGF-1R ECD revealed a significant clash between the vimentin head domain and IGF-1R (Extended Data Fig. 5b). This prompted us to investigate why only mssVIM, lacking the N-terminus, could bind to IGF-1R. By analyzing the staggered tetramer model of vimentin, the basic unit for vimentin filaments assembly^38^, we found that the predicted binding sites (319-339 aa) of vimentin to IGF-1R were occluded by the N-terminus of an adjacent vimentin molecule during tetramer formation (Fig. 5d). This suggests that full-length vimentin filaments mask the binding region for IGF-1R interaction. Furthermore, employing peptides of vimentin 71-90 aa, which mask the binding site on IGF-1R (319-339 aa), resulted in a restoration of MCF-7 cell migration in a dose-dependent manner when co-treated with the supernatant of WT THP-1 Mφ, whereas full-length recombinant vimentin failed to produce similar effects (Fig. 5e). Further experiments showed that the vimentin peptide 71-90 aa also inhibited in mammosphere invasion (Fig. 4f).

Pritumumab (PTB), a fully human IgG1 (kappa) monoclonal antibody originally isolated from a patient with cervical carcinoma, is currently in clinical trials for curing glioblastomas. Its functional epitope (288-368 aa) on vimentin’s rod region includes the predicted binding site of mssVIM to IGF-1R (Fig. 5d)^39,40^. PTB weakens the pro-migration ability of WT THP-1 Mφ supernatant in wound healing assay similarly to anti-C-VIM (Fig. 5g), which was further verified in the mammosphere invasion assay (Fig. 5h), indicating the potential clinical indication of mssVIM in anti-cancer therapy.

IGF-1 is the canonical ligand for IGF-1R. However, it was not detected in the mass spectrometry analysis we performed on the tumor interstitial fluid. We then ruled out the pro-tumor migration function of IGF-1 by the wound healing assay (Fig. 5g). Although local 3D Zernike descriptor-based protein docking analysis (LZerD) showed that the average affinity ranksum score for the top 50 poses of IGF-1R with mssVIM was relatively lower than that for IGF-1, the interface-tolerant score rank (ITScore) was comparable to that of IGF-1 (Extended Data Fig. 5c), indicating that the interaction between mssVIM and IGF-1R was relatively stable. Together, these results highlight mssVIM as a novel ligand-like molecule capable of activating IGF-1R with a comparable affinity but distinct cellular function compared to the canonical ligand IGF-1 (Extended Data Fig. 5d).

### Ribosomal S6 kinase (RSK) is activated upon mssVIM recognizing IGF-1R

IGF-1 typically activates the mitogen-activated protein kinase/mitogen-activated extracellular signal-regulated kinase (MAPK/MEK) and protein kinase B (AKT) signaling pathways upon binding to IGF-1R (Fig. 6a)^41^. Therefore, we investigated whether mssVIM also activates these signaling pathways. Western blots showed that treatment with WT THP-1 Mφ supernatant did not lead to significant changes in the activation levels of MEK, ERK, and AKT compared to treatment with VIM KO THP-1 Mφ supernatant (Fig. 6b). This suggests that mssVIM may not exert its effects through the canonical IGF-1 signaling pathways. To investigate the downstream signals activated by mssVIM, we employed a human phospho-kinase array, which simultaneously detects the phosphorylation of 37 kinases, including AKT and ERK (Extended Data Fig. 6a). The results further excluded the activation of the AKT and MEK signaling pathways. Intriguingly, ribosomal S6 kinase (RSK) was identified as the most significantly activated protein (Fig. 6c-e and Extended Data Fig. 6b), which was further confirmed by western blot analysis treated with the supernatants from WT and VIM KO THP-1 Mφ, respectively (Fig. 6f).

**Fig. 6.**
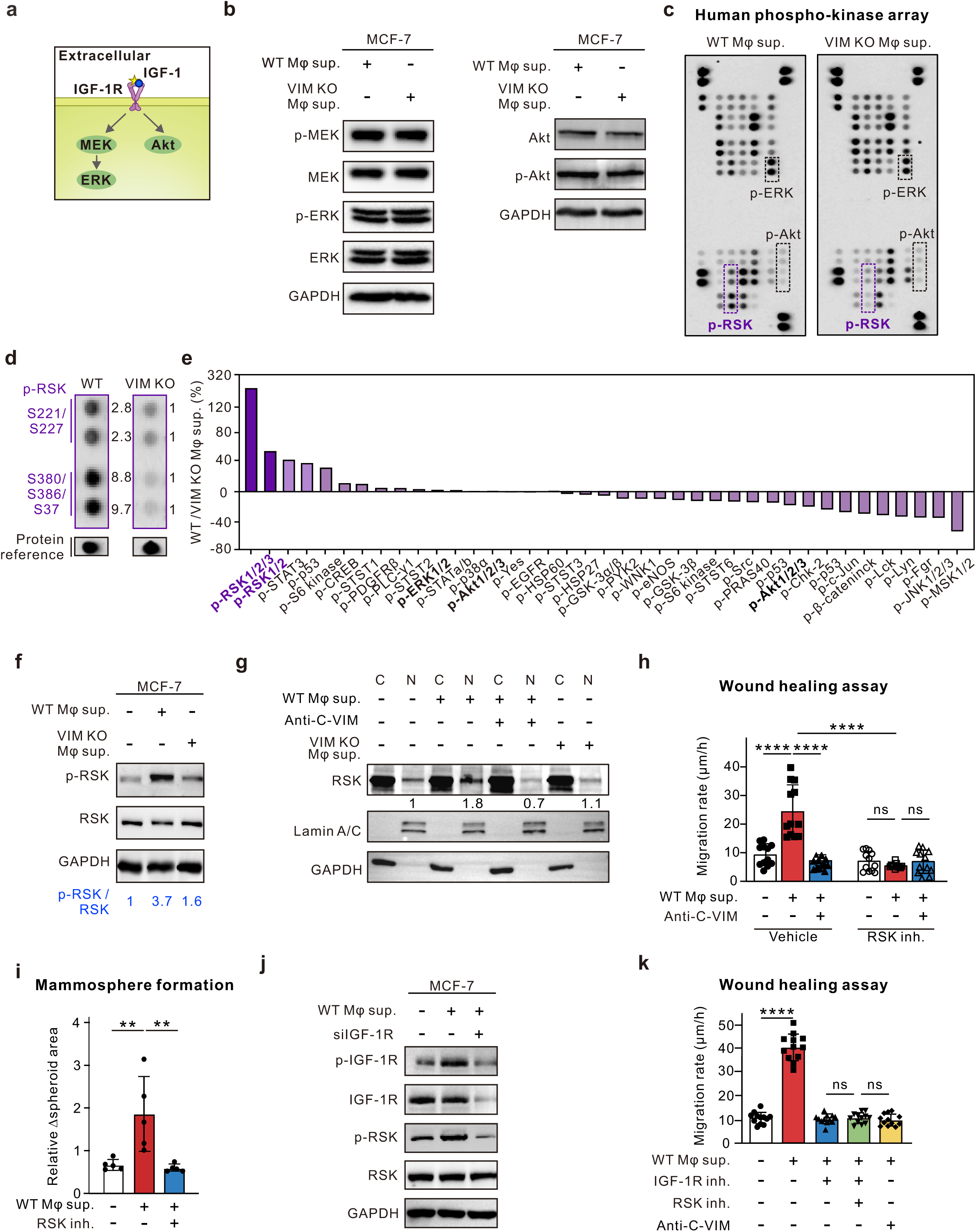
RSK in cancer cells is activated via mssVIM-IGF-1R signaling. **a**, Schematic diagram illustrating the activation of the IGF-1-IGF-1R downstream signaling pathway. **b**, Western blot analysis of p-MEK, MEK, p-ERK, and ERK levels (left panel) and p-AKT and AKT levels (right panel) in MCF-7 cells treated with the supernatants from WT or VIM KO THP-1 Mφ. GAPDH was used as a loading control. **c**, Human phospho-kinase array analysis of MCF-7 cells treated with the supernatants from WT or VIM KO THP-1 Mφ. p-ERK and p-Akt are highlighted with black boxes, and p-RSK with purple boxes. **d**, Comparison of p-RSK levels in MCF-7 cells treated with supernatants from WT or VIM KO THP-1 Mφ. The numbers indicate the intensity of the corresponding bands normalized to the VIM KO group. **e**, Quantification of all phospho-kinase levels in (**c**). **f**, Western blot analysis of p-RSK and RSK levels in MCF-7 cells treated with supernatants from WT or VIM KO THP-1 Mφ. GAPDH was used as a loading control. Ratios of p-RSK to RSK are indicated by blue numbers. **g**, Western blot analysis of RSK levels in the cytoplasmic and nuclear fractions of MCF-7 cells treated with the WT or VIM KO THP-1 Mφ supernatants, with or without anti-C-VIM. Lamin A/C and GAPDH serve as a nuclear and cytoplasmic controls, respectively. The numbers in the blots indicate RSK levels normalized to their corresponding nuclear levels. **h**, Quantification of the migration rate of MCF-7 cells treated with THP-1 Mφ supernatants, with or without anti-C-VIM or RSK inhibitor FMK in the wound healing assay. **i**, Quantification of fold changes in the mammosphere area treated with THP-1 Mφ supernatants, with or without RSK inhibitor FMK. **j**, Western blot analysis of p-IGF-1R, IGF-1R, p-RSK, and RSK levels in WT and siIGF-1R MCF-7 cells treated with THP-1 Mφ supernatants. GAPDH serves as a loading control. **k**, Quantification of the migration rate of MCF-7 cells treated with THP-1 Mφ supernatants, with or without anti-C-VIM, RSK, and IGF-1R inhibitors. Data are presented as mean ± S.D. from three independent experiments. ns *P* > 0.05; \*\*\**P* < 0.001, \*\*\*\**P* < 0.0001: one-way ANOVA multiple comparisons (**i**, **h**, **j**).

Previous studies have shown that upon activation, RSK typically translocates to the nucleus, where it activates transcription factors^42,43^. Cellular fractionation experiments demonstrated that treatment with WT THP-1 Mφ supernatant promoted the nuclear translocation of RSK (Fig. 6g), with nuclear RSK levels steadily increasing within the first 12 hours of stimulation (Extended Data Fig. 6c). The RSK inhibitor FMK significantly inhibited the pro-migration effect of mssVIM in both 2D wound healing assay and 3D mammosphere invasion assay (Fig. 6h, i and Extended Data Fig. 6d), highlighting the critical role of RSK in mssVIM-induced tumor migration.

To clarify the upstream and downstream relationship between RSK and IGF-1R, IGF-1R knockdown experiments were conducted, revealing a significant suppression of RSK activation (Fig. 6j). No additional suppression was observed when combined with WT THP-1 Mφ supernatant treatment, indicating that RSK is a key downstream signal regulated by IGF-1R upon mssVIM stimulation. Further combinatorial drug experiments confirmed this finding, showing that the IGF-1R inhibitor did not exert additional inhibitory effects when used in conjunction with the RSK inhibitor (Fig. 6k). Together, these results establish the central role of the IGF-1R-RSK axis in mssVIM-induced tumor migration.

### mssVIM enhances the level of integrin αVβ6 in tumor cells via the IGF-1R-RSK axis

The significantly enhanced migration of cancer cells stimulated by mssVIM suggests the upregulation of proteins involved in migration-related functions. RNA sequencing analysis revealed a significant number of candidate proteins associated with cell migration and adhesion following stimulation with the supernatants from WT but not VIM KO THP-1 Mφ (Fig. 7a). Focal adhesion (FA), a cellular component closely associated with cell migration^44,45^, was among the top upregulated cellular components (Fig. 7b). Immunofluorescence analysis revealed FA marker vinculin was significantly upregulated upon treatment with WT THP-1 Mφ supernatant compared to the mock group and the VIM KO THP-1 Mφ supernatant treatment group (Extended Data Fig. 7a-c). Among the upregulated focal adhesion-related genes, a pair of integrin genes, *ITGAV* and *ITGB6*, was found to be significantly increased (Fig. 7c)^46^. Previous research has demonstrated that the integrin αVβ6 dimer, encoded by *ITGAV* and *ITGB6*, plays a crucial role in the malignant behavior of cancer cells^46^. Quantitative PCR and western blots analysis further confirmed the upregulated RNA and protein levels of integrin αV and β6 upon stimulation with WT THP-1 Mφ supernatant (Fig. 7d, e).

**Fig. 7.**
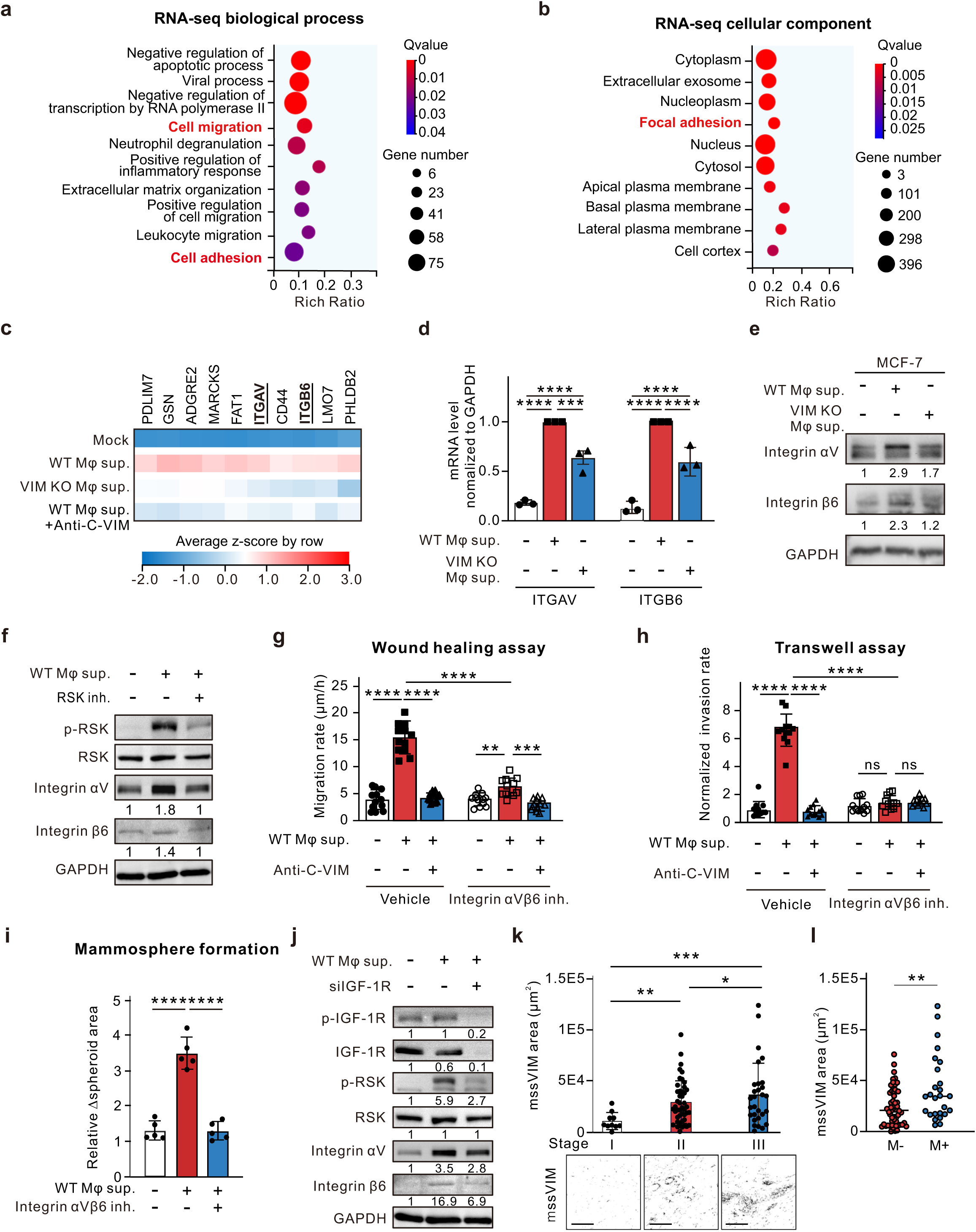
mssVIM enhances integrin expression to promote cancer cell migration via the mssVIM-IGF-1R-RSK signaling. **a**, **b**, Enrichment analysis of biological processes and cellular components from significantly upregulated genes following supernatant treatment of WT THP-1 Mφ by RNA-sequencing. The color of the bubbles displayed from red to blue indicates the descending order of − log10(Padj). The sizes of the bubbles are displayed from small to large in ascending order of gene counts. The x- and y-axes represent gene ratios and GO terms, respectively. **c**, Heatmap of significantly differentially expressed genes in MCF-7 cells treated with supernatants from indicated conditions (n = 3 independent experiments). **d**, Quantitative RT-PCR measurement of ITGAV and ITGB6 mRNA levels in MCF-7 cells treated with WT or VIM KO THP-1 Mφ supernatants. **e**, Western blot analysis of integrin αV and β6 expression in MCF-7 cells treated with WT or VIM KO THP-1 Mφ supernatants. GAPDH serves as a loading control. Numbers indicate levels of integrin αV and integrin β6 expression normalized to those of GAPDH and the control group. **f**, Western blot analysis of p-RSK, RSK, integrin αV and β6 levels in MCF-7 cells treated with THP-1 Mφ supernatant, with or without the RSK inhibitor FMK. GAPDH serves as a loading control. Numbers in the blots indicate the levels of integrin αV and integrin β6 expression normalized to those of GAPDH and the control group. **g**, **h**, Quantification of the mean migration rate (**h**) or the invasion rate (**i**) of MCF-7 cells treated with THP-1 Mφ supernatants, with or without anti-C-VIM or the integrin αVβ6 inhibitor EMD527040, in the wound healing assay and the transwell assay, respectively. **i**, Quantification of fold changes in the mammosphere area treated with THP-1 Mφ supernatants, with or without integrin αVβ6 inhibitor EMD527040. **j**, Western blot analysis of p-IGF-1R, IGF-1R, RSK, p-RSK, integrin αV and β6 levels in WT and siIGF-1R MCF-7 cells treated with WT THP-1 Mφ supernatant. GAPDH serves as a loading control. Numbers in the blots indicate the levels of integrin αV and β6 expression normalized to those of GAPDH and the control group. **k**, Quantification of mssVIM area among breast cancer patients from different stages. Scale bars, 100 μm. **l**, Quantitative analysis of mssVIM areas in relation to lymph node metastasis in breast cancer patients, lymph node metastasis is indicated as M+; absence of lymph node metastasis is indicated as M-. Data are presented as mean ± S.D. from three independent experiments. ns *P* > 0.05; \*\*\**P* < 0.001, \*\*\*\**P* < 0.0001: one-way ANOVA multiple comparisons (**d**, **g**, **h**, **i**).

Consistent with previous studies indicating that nuclear translocation of RSK promotes integrin transcription^47^, RSK inhibition significantly suppressed integrin αVβ6 upregulation (Fig. 7f). Additionally, both the pan-integrin inhibitor GRGDSP and the specific αVβ6 integrin inhibitor EMD527040 diminished the pro-migration effect induced by the WT THP-1 Mφ supernatant (Fig. 7g, h, and Extended Data Fig. 7d-f). The *in vivo*-like mammosphere invasion assay further confirmed that the integrin inhibitor reduced tumor invasion levels (Fig. 7i). Knockdown of IGF-1R impaired the downstream activation of RSK and subsequent expression of integrin αVβ6 (Fig. 7j). We hence propose that integrin αVβ6 plays a pivotal role in mssVIM-dependent cancer cell migration. Taken together, the activation of the IGF-1R-RSK axis by mssVIM enhances the transcription of *ITGAV* and *ITGB6*, leading to increased expression of integrin αVβ6 on the tumor cell surface, ultimately promoting tumor migration.

To investigate whether the level of mssVIM is quantitatively correlates with the extent of malignancy, we performed immunofluorescence staining on total 60 breast cancer patient tissue samples with increased clinical staging. The anti-N-VIM and anti-C-VIM antibodies were co-stained to distinguish mssVIM as the un-overlapped anti-C-VIM signal (Extended Data Fig. 7g). We found that the level of mssVIM is positively correlated with tumor malignancy by quantifying its level in progressive clinical stages from I to III (Fig. 7k). Further, we analyzed the correlation of lymph node metastasis and mssVIM levels within tumor tissues, and found that mssVIM levels were significantly higher in the group with lymph node metastasis (Fig. 7l). These findings provide new evidence for mssVIM as a potential tumor biomarker and therapeutic target.

## Discussion

The co-evolution of malignant cells with tumor microenvironment (TME) is a key driver of tumor metastasis^48^. TAMs are recognized as major contributors^28,49,50^, with their secretory profile playing a crucial role in tumor progression^51^. In this study, we identified a novel variant of the cytoskeletal protein vimentin, mssVIM, which is selectively secreted by TAMs and serves like a ligand to trigger a signaling cascade that directly enhances tumor migration and metastasis (Fig. 8). This discovery not only deepens our understanding of TAM-mediated tumor metastasis but also suggests that targeting the mssVIM cascade we revealed here may offer a novel therapeutic approach for cancer treatments.

**Fig. 8.**
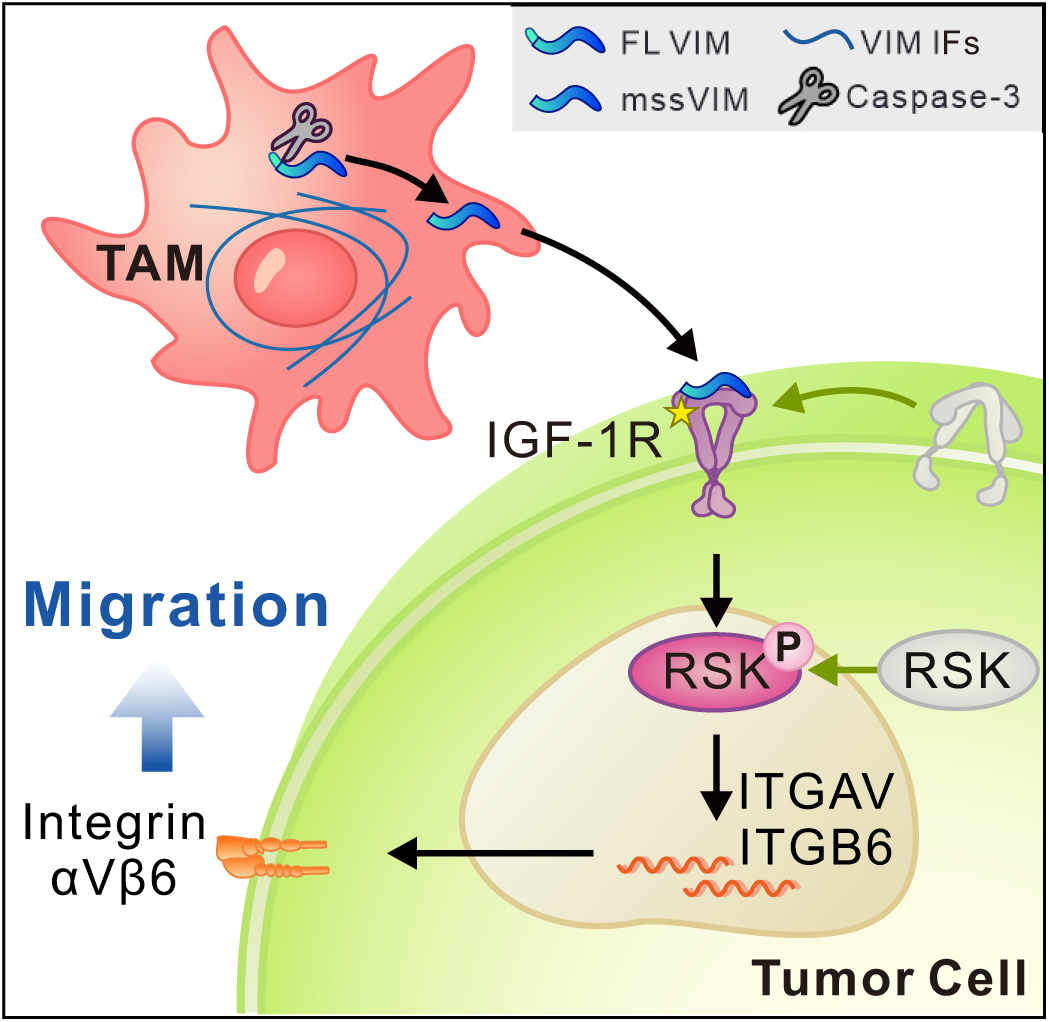
The schematic diagram of the mssVIM directed cancer cell migration by engaging IGF-1R.

Vimentin, traditionally recognized for its mechanical functions as a cytoskeletal protein, has recently gained considerable attention for its non-mechanical roles in cellular processes, particularly in inflammation and pathogen infection^52^. It acts as a scaffold for pattern recognition receptors (PRRs) during inflammatory responses and facilitates SARS-CoV-2 invasion by acting as an attachment factor^53–55^. During Zika virus infection, vimentin supports viral replication by interacting with ribosome receptor binding protein 1 (RRBP1)^56^. Despite of these recognized non-cytoskeletal functions, our study, for the first time, reveals the existence of a cleaved variant of vimentin and elaborates on its ligand-mimicking role in the context of cancer.

Previous studies show that extracellular vimentin typically exists in full-length form^24,26,55,57–64^ and is primarily released through exosomes to transport between cells^26,65^. Vimentin-exosomes released by adipocytes protect fibroblasts from osmotic stress, thereby promoting wound healing^65^. Vimentin-exosomes secreted by astrocytes enhance C3bot binding to neuronal membranes, thereby affecting the functions of astrocytes and neurons^26^. Different from the full-length extracellular vimentin reported in all early studies, mssVIM we found is cleaved and secreted via the type I unconventional secretion system, indicating that mssVIM and full-length secreted vimentin have completely distinct processing mechanisms within the cytoplasm, corresponding to their unique functions.

Full-length vimentin cannot replicate the pro-migration function of mssVIM, suggesting potential configurational distinctions in this variant. Indeed, the lack of the N-terminal head domain disables vimentin from assembling to filaments by disrupting the staggered tetramer formation, based on both our data and a previous study^66^. Once vimentin assembles into filaments, the polymeric structure masks its binding sites on the head domain for IGF-1R. Removal of the head domain as mssVIM form exposes the sites for the interaction. Thus, a detailed understanding of the molecular configuration and processing of mssVIM is vital for elucidating its role in cancer progression and its potential as a therapeutic target.

Various components within TME intricately interact to create a nurturing niche that facilitates tumor progression^67,68^. TAMs have been reported in pro-tumorigenic processes, including establishing an immunosuppressive environment through the expression of PD-L1, remodeling the extracellular matrix by secreting matrix metalloproteinases (MMPs), and promoting drug resistance by releasing cytokines such as IL-6 and IL-10^7,69^. Beyond these, our study emphasizes the uniqueness of TAMs by demonstrating that they exclusively secrete mssVIM to selectively and directly accelerate cancer cell migration capability, a phenomenon not detected in T cells, B cells, neutrophils, nor tumor cells.

In breast cancer treatment, where approximately 70% of cases express estrogen and progesterone receptors, traditional hormone therapy targeting the estrogen receptor (ER) pathway, including medications like tamoxifen and fulvestrant, encounter significant stumbling blocks due to the high incidence of both *de novo* and acquired resistance during therapy^70^. Despite advances in understanding resistance mechanisms, recurrence remains a critical issue^71,72^. The cue points underlying this issue may stem from the variability in TME cell-cell communication, including the effects of TAMs on tumor cells^73^. In this context, our study introduces a critical yet previously overlooked factor, mssVIM. This unique contribution of TAMs thus plays a pivotal role in driving breast cancer cell migration and metastasis. The clinical phase II drug pritumumab, which targets vimentin, inhibits the pro-migratory function of mssVIM, suggesting its potential application in anti-tumor therapy.

In certain contexts, the MCF-7 mammosphere invasion model and the mouse orthotopic injection model we used in this study effectively simulate the initiation of ductal carcinoma *in situ* (DCIS), a noninvasive form of breast cancer^74,75^. Clinically, approximately 30% of DCIS cases progress to invasive ductal carcinoma (IDC), while many patients receive overtreatment like surgery and radiotherapy^76,77^. This underscores the need for a more appropriate patient stratification standard to distinguish invasion-prone DCIS from noninvasive DCIS^78^. High macrophage infiltration has always been a TME signature correlated with poorer prognostic outcomes^78^. Our study extends this acknowledgment by identifying TAM’s unique secretion factor mssVIM and specific downstream IGF-1R-RSK signaling. The ultimately increased integrin αvβ6 is significantly associated with cancer progression and recurrence^79^. Together, these findings suggest that targeting factors along this cascade potentiate a way to optimize patient stratification to reduce the risk of DCIS overtreatment, offering a more tailored approach to therapy.

Notably, our experiments demonstrated that mssVIM treatment not only elevated migration in breast cancer cells (MCF-7) but also in osteosarcoma (U2OS) and cervical cancer cells (HeLa). Cancer cell lines with low IGF-1R expression did not exhibit similar responses, nor did studies involving the reduction or blocking of IGF-1R expression in MCF-7 cells (Figure 4E, 4F, and 4H). These findings suggest that migration induced by mssVIM is largely dependent on IGF-1R expression in recipient tumor cells, which is recognized as a common oncoprotein in multiple cancer lineages^80^. This provides a strong rationale for future clinical assessments of anti-cancer therapies targeting mssVIM, which could offer new avenues for the treatment across various cancer lineages.

Taken together, our study revealed a novel mechanism by which TAMs contribute to breast cancer progression through the secretion of a cleaved variant of vimentin. This molecule acts as a signal mediator, promoting cancer cell migration and metastasis via the IGF-1R-RSK pathway. The discovery of mssVIM as a critical factor in TAM-driven tumorigenesis highlights the complexity of TME and underscores the importance of targeting TME components to overcome therapeutic resistance. Importantly, the broad-spectrum impact of mssVIM on different cancer types suggests that therapies targeting this pathway may have far-reaching implications beyond breast cancer. These findings open new avenues for developing mssVIM-IGF-1R-focused therapies, offering potential strategies to improve clinical outcomes in cancer patients.

## Methods

### Ethics statement

This study has been approved by the Institutional Review Board of Shanghai East Hospital (China) (No. 2024YS-071). All samples were collected after obtaining written informed consent from patients. All animal experiments were conducted following institutional ethics requirements under the animal user permit (No. A2024008) approved by the Animal Care and Use Committee of the Shanghai Institute of Immunity and Infection.

### Breast cancer sample collection and preparation

Tissue samples were obtained from breast cancer patients diagnosed with stage III ductal carcinoma of the breast, characterized by lymphovascular invasion. The informed consent in accordance with the guidelines approved by the Ethics Committee of Shanghai East Hospital. Post-collection, the tissue samples were gently rinsed with phosphate-buffered saline (PBS) and preserved in the transfer medium consisting of PBS and 1% penicillin-streptomycin (P/S) before isolation.

### (Peri)tumor interstitial fluid collection

Tumor and peri-tumor tissues were finely minced using a surgical scalpel. The minced tissues were subsequently submerged in PBS containing 1% P/S and incubated at 37°C with 5% CO_2_ for 1 h. The supernatant was collected and concentrated using Amicon Ultra Centrifugal Filters, producing the peritumoral interstitial fluid.

### Primary macrophages isolation

Dissociation and wash buffer were prepared prior to tumor digestion. The dissociation buffer comprised 4 mg/mL Collagenase Type IV (Stemcell, 07909), 1 mg/mL hyaluronidase (Stemcell, 07461), and 10 U/µL DNase I (Beyotime, D7073). This mixture was prepared in Human Mammary Epithelial Cell Basal Medium (Gibco, M171500) supplemented with Mammary Epithelial Growth Supplement (MEGS, Gibco, S0155) and 1% P/S, and then filtered with 0.22 µm polyether sulfone (PES) syringe filter. The wash buffer consisted of PBS containing 2% FBS and 1% P/S.

Minced tumor tissue was digested with dissociation buffer at 37°C on a shaker for 45 min. The cells were centrifuged at 300×g for 10 min at room temperature (RT). The supernatant was discarded and the pellet was resuspended in wash buffer. Macrophages were separated from the supernatant with another centrifugation for 10 min, washed with wash buffer and further cultured in RPMI-1640 supplemented with 10% Fetal Bovine Serum (FBS), 2 mM L-glutamine (Gibco, 35050061) and 50 ng/ml M-CSF (Sino biologcal, 11792-HNAH) at 37°C with 5% CO_2_.

### Primary breast cancer cell isolation

Primary breast cancer cells were separated using a modified sedimentation technique. Briefly, after the supernatant was removed for macrophage isolation, the remaining solution was allowed to settle at RT for 20 min to facilitate the sedimentation of cancer cells. Then, half of the volume above the pellet was gently aspirated, and an equal volume of fresh wash buffer was added. These steps were repeated for ten times, and the cells were subsequently pelleted by centrifugation at 300×g for 10 min. The final pellet, enriched in primary breast cancer cells, was collected and processed for further analysis.

### Cell culture

The human leukemia monocytic cell line THP-1 was cultured in Roswell Park Memorial Institute medium (RPMI) 1640 (Gibco, 11875093) supplemented with 10% fetal bovine serum (FBS) (Gibco, 10091148), 1% penicillin and streptomycin (P/S) (Gibco, 15140122), 50 μg/mL 2-mercaptoethanol (Sigma, M6250), 1% non-essential amino acids (Gibco, 11140050) and 1% sodium pyruvate (Gibco, 11360070) at 37°C, 5% CO_2_. MCF-7, MCF-7-luciferase (MCF-7-LUC), HeLa, U2OS and HUVEC cell lines were cultured in Dulbecco’s Modified Eagle Media (DMEM) (Biological Industries, 01-055-1A) supplemented with 10% FBS, 1% P/S at 37°C, 5% CO_2_. U937, HL-60, Jurkat and Raji cells were cultured in RPMI-1640 supplemented with 10% FBS, 1% P/S at 37°C, 5% CO_2_. TMED10 KO THP-1 cell line is kindly provided by Dr. Liang Ge (Tsinghua University, Beijing 100084, China).

### Macrophages differentiation and supernatant collection

The THP-1 monocytes differentiated into macrophage-like cells by treatment with 100 ng/mL phorbol myristate acetate (PMA) for 48 h. Then, the culture medium was changed to RPMI-1640 supplemented with 10% FBS and 1% P/S and incubated for additional 24 h. The supernatant was then carefully collected and centrifuged at 3000×g for 30 min. After centrifugation, the supernatant was subsequently filtered through 0.22 µm PES syringe filters and prepared for further applications.

### Tissue staining and immunofluorescence

Paraffin-embedded sections of breast cancer tissue microarray were acquired from Biossci (Shanghai, China). Initially, the tissue sections were deparaffinized in xylene and subsequent rehydration through a graded series of ethanol solutions. For antigen retrieval, the sections were heated in a citrate buffer. To prevent non-specific binding, the sections were then blocked with 10% goat serum at RT for 30 min. Subsequently, the sections were incubated overnight at 4°C with primary antibodies including collagen IV antibody (dilution 1:400, Abcam, ab214417), CD68 antibody (Maxim, Kit-0026), vimentin mouse antibody (anti-C-VIM, dilution 1:40, Sigma, V6630), and pan-cytokeratin (pan-CK) antibody (dilution 1:800, ZSGB-BIO, ZM-0069). After thorough washing with phosphate-buffered saline (PBS), the sections were incubated with secondary antibodies, Goat Anti-Rabbit IgG (HRP, 1:4000, Abcam, ab205718) and Goat Anti-Mouse IgG (HRP, 1:2000, Abcam, ab205719), for 45 min at 37°C. Following this, the sections were stained with fluorescent dyes: FITC (dilution 1:300, AAT Bioquest, 11060), CY3 (dilution 1:600, AAT Bioquest, 11065), CY5 (dilution 1:400, AAT Bioquest, 11066), and 430 (dilution 1:400, AAT Bioquest, 45096) for 10 min at RT. Finally, the sections were counterstained with DAPI to visualize the nuclei and mounted with an anti-fade mounting medium. Semi-quantification was performed using Fiji software (National Institutes of Health, version 1.53f). Tissue microarray was obtained from Bioaiteh (Shanxi, China). Staining was performed by Servicebio (Wuhan, China). Primary antibodies including vimentin mouse antibody (anti-C-VIM, dilution 1:40, Sigma, V6630) and vimentin rabbit antibody (anti-N-VIM, dilution 1:200, Cell Signaling Technology, 5741).

### Western blots

Cells were lysed in RIPA lysis buffer (Beyotime, P0013B) supplemented with 50 × protease and phosphatase inhibitor (Beyotime, P1045). After centrifugation at 800×g for 5 min, the supernatant was transferred to a new tube. Protein concentrations were determined using the Enhanced BCA Protein Assay Kit (Beyotime, P0010) and adjusted to the same level. The lysates were mixed with 6× SDS sample buffer (Tanon, 180-8201D) and separated by 10% SDS-PAGE gels. Proteins were transferred onto a polyvinylidene fluoride (PVDF) membrane in Tris-buffered saline with 0.1% Tween-20 (TBST). The membrane was then blocked with 5% non-fat milk (BD Difco, 8011939) for 30 min at RT. Primary antibodies were applied at dilutions according to the manufacturer’s instructions and incubated overnight at 4°C. Horseradish peroxidase (HRP)-linked secondary antibodies were diluted according to manufacturer’s protocols and incubated with 1 h at RT. Proteins were detected from the membrane with Western blots ECL (Tanon, 180-501). Band intensities of detected protein were quantified using ImageJ software (National Institutes of Health, version 1.53f) and normalized to GAPDH levels as a loading control. The following antibodies were used: GAPDH rabbit antibody (dilution 1:5000, Sigma, G9545); vimentin mouse antibody (anti-C-VIM, dilution 1:1000, Sigma, V6630); vimentin (D21H3) rabbit antibody (anti-N-VIM, dilution 1:1000, Cell Signaling Technology, 5741); vimentin chicken antibody (anti-FL-VIM, dilution 1:1000, Abcam, ab24525); IGF-1R rabbit antibody (dilution 1:1000, Cell Signaling Technology, 3918); p-IGF-1R rabbit antibody (dilution 1:1000, Cell Signaling Technology, 3024); His-Tag mouse antibody (dilution 1:1000, Proteintech, 66005-1-lg), Akt1/2/3 rabbit antibody (dilution 1:1000, Abmart, T55561), p-Akt rabbit antibody (dilution 1:1000, T40067), MEK1/2 mouse antibody (dilution 1:1000, Cell Signaling Technology, 4694), p-MEK1/2 rabbit antibody (dilution 1:1000, Cell Signaling Technology, 9154), ERK1/2 rabbit antibody (dilution 1:1000, Cell Signaling Technology, 4695); p-ERK1/2 rabbit antibody (dilution 1:1000, Cell signaling Technology, 4370); RSK rabbit antibody (dilution 1:1000, Cell Signaling Technology, 9355); p-RSK rabbit antibody (dilution 1:1000, Cell Signaling Technology, 9341); lamin A/C (dilution 1:1000, Cell Signaling Technology, 2032); integrin αV (dilution 1:1000, Millipore, MAB1953); integrin β6 mouse antibody (dilution 1:1000, Millipore, MAB2075Z); HRP-linked anti-mouse IgG antibody (dilution 1:5000, Cell Signaling Technology, 7076V); HRP-linked anti-rabbit IgG antibody (dilution 1:5000, Cell Signaling Technology, 7074V).

### Immunoprecipitation and co-immunoprecipitation

For immunoprecipitation, harvested supernatant from 6×10^5^ THP-1 macrophage was incubated with 10 µg of vimentin rabbit antibody (Abcam, ab137321) for 2 h at 4°C with rotation. Then, 20 µL of protein A/G agarose beads (Santa Cruz Biotechnology, SC-2003) were added to the sample and rotated at 4°C overnight. Following incubation, the beads were washed four times with wash buffer (50mM Tris-HCl pH 6.8, 100mM NaCl, 1mM EDTA). After the final wash, the beads were resuspended in 40 µL of 5× SDS sample buffer and boiled for 10 min to elute the bound proteins. The immunoprecipitated proteins were then loaded onto SDS-PAGE gels, and subjected to Western blot analysis.

For co-immunoprecipitation, harvested supernatant from 1 × 10^7^ THP-1 macrophages were pre-incubated with 10 µg of human recombinant IGF-1R (Sino biological, 10164-H08H) at RT for 30 min. The procedure was then carried out as described for immunoprecipitation, except for the initial antibody incubation step.

### Cell fractionation analysis

For cell membrane protein extraction, MCF-7 cells treated with THP-1 supernatant were collected and processed by Subcellular Protein Fractionation Kit for Cultured Cells (Thermo Scientific, 78840) according to the manufacturer’s instructions. Subsequently, proteins were immunoprecipitated with the phospho-tyrosine mouse antibody (dilution 1:100, Cell Signaling Technology, 9411) and analyzed by mass spectrometry.

### Mass spectrometry

Protein bands of interest were excised from SDS-PAGE gels and destained using a solution containing 25 mM NH_4_HCO_3_ and acetonitrile (ACN) (1:1, v/v) at 37°C for 10 min. The gel pieces were then subjected to reduction with 10 mM dithiothreitol (DTT) at 56°C for 1 h, followed by alkylation with 50 mM iodoacetamide in the dark at RT for 45 min. Proteins were precipitated by adding ice-cold acetone and incubated at −80°C for 30 min, then at −20°C for additional 30 min. After centrifugation at 20,000×g for 5 min, the supernatant was discarded, and the protein pellet was washed with 1 mL of ice-cold acetone. The pellet was resuspended in 50 mM NH_4_HCO_3_ to a final protein concentration of 5 mg/mL. In-gel tryptic digestion was performed overnight at 37°C with a 1:50 (w/w) trypsin-to-protein ratio using MS Grade Trypsin Protease (Thermo Scientific, 90057). The digested peptides were acidified with 1% trifluoroacetic acid (TFA) and centrifuged at 5000×g for 5 min. The supernatant containing the peptides was aliquoted and stored at −80°C until analysis by mass spectrometry analysis using a Q Exactive Plus and Vanquish Flex (Thermo Scientific).

Raw nano-LC−MS/MS files were analyzed using Proteome Discoverer 2.4 software (Thermo Scientific) with the Sequest HT search engine. A composite FASTA file containing Homo sapiens proteins from UniProt was used for database searching and protein identification. The data were analyzed with the following parameters: trypsin as the enzyme, which digests proteins at arginine (R) and lysine (K) residues, fixed carbamidomethyl modification of cysteine residues, variable oxidation of methionine, a maximum of two missed cleavages, 5 ppm precursor tolerance, and 0.02 Da MS/MS tolerance. Data were filtered at 1% peptide FDR using the percolator node. The ‘precursor ions quantifier node’ was used for quantification by peak area in fold-change analysis.

### Phosphopeptide enrichment and mass spectrometry

Pierce^TM^ TiO_2_-coated magnetic beads (Thermo Scientific, 88811) were used for phosphopeptide enrichment as the manufacturer’s protocol. The lyophilized peptide mixture was resuspended in 200 μl 80% (vol/vol) ACN with 2% (vol/vol) formic acid before incubation with 10 μl TiO_2_ magnetic beads for 1 min. Subsequently, the tubes were placed on a magnetic plate for 1 min, the supernatant was removed, and the beads were washed three times with 200 μl binding buffer provided in the kit. Phosphopeptides were eluted for 10 min with 30 μl elution buffer provided in the kit and lyophilized to dryness before liquid chromatography-mass spectrometry analysis.

### RNA sequencing and transcriptome analysis

Total RNA was extracted from THP-1 and MCF-7 cells using TRIzol™ reagent (Invitrogen, 15596026) according to the manufacturer’s instructions. RNA libraries were prepared following the mRNA Library Preparation (DNBSEQ) BGI-NGS-JK-RNA-001 A0 protocol. Raw sequencing data were processed with SOAPnuke to remove reads containing sequencing adapters^81^, low-quality bases (quality score ≤ 15) exceeding 20% of the read length, and reads with more than 5% unknown bases (‘N’). Clean reads were stored in FASTQ format.

Subsequent analysis was performed using the Dr. Tom Multi-Omics Data Mining System (https://biosys.bgi.com). Alignment and quantification of clean reads in the reference transcriptome were performed using Bowtie2. Gene expression levels were calculated with RSEM (v1.3.1)^82^. Differential expression was analyzed with DESeq2 (v1.4.5) using a |log2FoldChange| > 0.5 and Q Value ≤ 0.05 (or FDR ≤0.001)^83^. Heatmaps for visualizing gene expression differences were generated using pheatmap (v1.0.8).

Isoforms of THP-1 identified by Iso-seq were visualized using Integrative Genomics Viewer (IGV). To interpret the biological significance of differentially expressed genes, Gene Ontology (GO) (http://www.geneontology.org/) and Kyoto Encyclopedia of Genes and Genomes (KEGG) enrichment analyses were performed using Phyper. (https://en.wikipedia.org/wiki/Hypergeometric_distribution) based on the hypergeometric test. Statistical significance was determined by Q value adjustment with a threshold of Q Value ≤ 0.05.

### Protein purification

Vimentin was purified from BL21(DE3) pLysS cells carrying pET28a. 2 L cell culture was grown in LB medium supplemented with kanamycin and chloramphenicol at 37°C to OD600 = 0.8, then induced with 1 mM IPTG and incubated at 16°C for 18 h. Cells were harvested and resuspended in lysis buffer (50 mM Tris-HCl pH 8.0, 25% sucrose, 1 mM EDTA, and 3 mM PMSF) and lysed by sonication on ice. Cell lysate was cleared by centrifugation at 15,000×g for 30 min at 4°C, and the pellet was collected. The pellet was washed, dissolved, and purified following the protocol described by Harald Herrmann^84^. The purified vimentin was renatured and concentrated to 2 mg/mL using an Amicon Ultra 30 kDa MWCO concentrator. The product was stored in aliquots at −80°C.

Caspase-3 was expressed in BL21(DE3) pLysS cells carrying pET22b. 2 L cell culture was grown in LB medium supplemented with kanamycin and chloramphenicol at 37°C to OD600 = 0.8, induced with 0.5 mM IPTG and incubated at 16°C for 16 h. Cells were harvested and resuspended in lysis buffer (25 mM HEPES pH 7.5, 150 mM NaCl, 5% glycerol, 1 mM PMSF) and lysed by sonication on ice. The cell lysate was cleared by centrifugation at 15,000×g for 40 min at 4°C, and the supernatant was loaded onto a 5 mL Ni-NTA column (Smart-life sciences, SA005500). The column was washed and eluted with elution buffer (25 mM HEPES, 150 mM NaCl, 10 mM β-mercaptoethanol, 5% glycerol, 250 mM imidazole). The eluted fractions were concentrated and further purified using a 24 mL Superdex 200 10/300 GL column (GE Healthcare, 28990944) equilibrated with 20 mM HEPES, and 150 mM NaCl. The fractions containing target proteins were concentrated to 2 mg/mL, and stored at −80°C.

### *In vitro* vimentin filaments assembly

Equal volumes of purified vimentin (2 mg/mL) and assembly buffer (45 mM Tris-HCL pH7.0, 100 mM NaCl) were gently mixed and incubated at 37°C for 1 h in a water bath to facilitate filament formation. Assembly was stopped by the addition of an equal volume of stop buffer (25 mM Tris-HCL pH 7.0, 50 mM NaCl, 0.2% glutaraldehyde) to the reaction mixture on ice.

### *In vitro* caspase-cleavage assay

Purified vimentin (2 mg/mL) was incubated with caspase-3 (2 mg/mL) at a molar ratio of 1:2 at 37°C for 1 h. The reaction was stopped by the addition of an SDS loading buffer. The samples were analyzed by SDS-PAGE to assess the extent of vimentin cleavage.

### Negative Stain Electron Microscopy (NSEM)

For NSEM grid preparation, 5 μL aliquots of the fixed sample were applied onto a carbon-coated electron microscopy grid that had been previously glow-discharged. The grids were incubated at RT for 1 min to allow adsorption of the sample. Excess fluid was removed, and 2% (w/v) uranyl acetate in water was then applied for 30 s to provide contrast. The grids were then air-dried and observed in a Tecnai G2 F20 TWIN microscope (FEI, Thermo Fisher Scientific).

### Mouse model experiments

For intravenous injection (I.V.) in the tail vein, female NOG mice aged 5 to 6 weeks were selected and divided into three groups. 5×10^5^ MCF-7 cells, treated with or without macrophage supernatant were injected into the lateral tail vein. RPMI-1640 or supernatant was administered every 5 days via tail vein injection. At 15 days post-inoculation, bioluminescent imaging was conducted by intraperitoneally injecting each mouse with 0.2 mL of a 15 mg/mL D-Luciferin (MedChemExpress, HY-12591), and visualized using the IVIS 200 (Xenogen, PerkinElmer) imaging system, with data analyzed using Living Image® (version 4.4).

For orthotopic injection (O.I.) on mammary fat pad, female nude mice, aged 6 to 7 weeks, were selected. 4×10^6^ MCF-7 cells, treated with or without macrophage supernatant, were mixed with matrigel (1:1, BD Biosciences) and then injected into the inguinal mammary fat pad. RPMI-1640 or supernatant was administered every 5 days via *in situ* or intratumoral injection, starting 10 days after the initial injection.

For immunohistochemistry analysis, harvested mouse tumors and lungs were immersion fixed in 10% neutral buffered formalin for 24 h and embedded in paraffin. Sections (4 μm thick) were mounted on silanized slides (Dako), deparaffinized, stained with hematoxylin and eosin (H&E), dehydrated, cleared in xylene, and mounted with neutral balsam for microscopic analysis. Detailed analysis was performed with Halo (indica labs, v3.6.4134).

### Vimentin CRISPR knockout cell line generation

Vimentin-knockout cells were generated using CRISPR/Cas9 system based on pSpCas9 (BB)-2A-GFP vector (Addgene, 48138) with two targets as previously described^85^. Primers for vimentin target 1 were 5’-CACCGTGGACGTAGTCACGTAGCTC-3’ and 5’-AAACGAGCTACGTGACTACGTCCAC-3’. Primers for vimentin target 2 were 5’-CACCGCAACGACAAAGCCCGCGTCG-3’ and 5’- AAACCGACGCGGGCTTTGTCGTTGC-3’. Transfected cells were detached at 24 h post-transfection and sorted with FACS Aria II (BD Biosciences) using low-intensity GFP-expression pass gating, and then cells were plated onto 96-well plates supplemented with DMEM containing 20% FBS and 10 mM HEPES with a single cell per well. CRISPR clones were cultivated for two weeks before selecting clones with no discernible target protein expression by Western blots.

### Random migration assay

Cells were collected from culture dishes using trypsin-EDTA and resuspended in 24-well plates at a density of 2×10^4^ cells/well. Cells were allowed to adhere and spread overnight. Before imaging, cells were incubated with Hoechst 33342 (MedChemExpress, HY-15559) for 10 min and washed twice with PBS, culture medium was then added to each well. Imaging was performed using Olympus IX73 inverted microscope with a UplanFL 10×/0.3 objective lens and controlled by CellSens Dimension software (Olympus, version 1.18). Average migration velocity and directional migration duration were quantified by tracking nuclear movement using Imaris Spots plugins (Bitplane). The recording was set as every 6 min for 12 h. Only cells that did not collide with one another were selected for measurement. The parameters for analysis in Imaris v9.2 (Bitplane) were configured with a 15 µm estimated xy diameter, a maximum distance of 30 µm, and a maximum gap size of 3 frames.

### Wound healing assay

Cells were seeded in a 6-well cell culture plate with a density of 2.5×10^5^/cm^2^ and incubated at 37°C with 5% CO_2_ overnight. Subsequently, the cell monolayers were scratched with a sterile pipette tip to create linear wounds and washed with PBS to remove detached cells. Cells were incubated in the corresponding medium and observed using an Olympus IX73 inverted microscope with a UplanFL 4/0.13 objective (Olympus) for 15 h. The wound areas were quantified at each time point using ImageJ software (National Institutes of Health, version 1.53f). The cell migration rate (V) was calculated using the formula 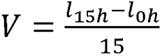, where V represents the migration rate, and l denotes the wound length.

### Transwell assay

2×10^5^ cells were added to the upper layer of the transwell chamber (Corning, 3470), and a corresponding clean RPMI-1640 medium or macrophage supernatant was added to the lower layer according to the experimental purpose. After 24 h culture, cells were fixed for 15 min using 4% paraformaldehyde in PBS, washed twice in PBS, stained with DAPI, and then washed twice more in PBS. After removing the upper cells with cotton swabs, the lower cells passing through the membrane were counted by the number of nuclei.

### Mammosphere formation assay

The MCF-7 cells were seeded in triplicate at a density of 8×10^4^ cells per well in 96-well spheroid microplates (Corning, CLS-AN-308). After five days, non-adherent compact spheroids were collected and gently washed with PBS. The spheroids were then transferred to matrigel-coated (Corning, 356234) 6-well plates. Once the matrigel solidified, different cell culture media were added according to the specific treatment conditions. The growth of mammospheres was monitored daily for four days.

### IGF-1R knockdown and IGF-1R multi-point mutation cell line generation

The small interference RNA (siRNA) against *IGF1R* was chemically synthesized by GenePharma (Shanghai, China) with the following sequence: 5’-CGACUAUCAGCAGCUGAAGTT-3’ and 5’-CUUCAGCUGCUGAUAGUCGTT-3’. The IGF-1R Mut plasmid was chemically synthesized by Sangon Biotech (Shanghai, China). The construct against IGF1R mRNA was referred to as IGF-1R siRNA and non-specific siRNA was referred to as siRNA negative control (siNC). MCF-7 cells grown to 70% confluence were transfected with IGF1R_pLX307 plasmid (Addgene, 98344) or IGF-1R siRNA or IGF-1R Mut plasmid using Lipofectamine 3000 transfection reagent (Invitrogen, L3000075) or Lipofectamine RNAiMAX reagent (Invitrogen, 13778075) following the manufacturer’s instructions. Cells were harvested after 72 h of transfection. The efficiency of IGF-1R knockdown was confirmed by Western blots.

### Alphafold prediction and the molecular docking

The structure of head-less vimentin and tumor-associated receptors was predicted using AlphaFold3. The predicted structure was then analyzed using PRODIGY v2.0 to obtain information on the binding affinity and dissociation constant, among other parameters.

The protein structures of mssVIM and extracellular domain (ECD) of IGF-1R were predicted using AlphaFold-Multimer following the protocol previously described^86^. Complex structures of mssVIM-IGF-1R ECD and mssVIM-FL-IGF-1R were also generated using AlphaFold-Multimer. The quality of the predicted complex structures was assessed based on their predicted TM-score (pTM) and interface pTM (ipTM), with the top-scoring pose chosen as the final conformation.

The molecular docking of mssVIM with IGF-1R was performed using the LZerD Web Server (version 2021.6) (https://lzerd.kiharalab.org/upload/upload/). IGF-1R was set as the receptor, while mssVIM and IGF-1 were set as ligands. Docking quality was evaluated using ITScore^87–89^. Differences in docking quality indicators between the two groups were analyzed using the independent two-sample *t*-test.

### Immunofluorescence microscopy

Cells were fixed with 4% paraformaldehyde (PFA) for 10 min, permeabilized with 0.1% Triton X-100 in PBS for 5 min, and then blocked with 5% BSA for 20 min. Primary and secondary antibodies were diluted in 5% BSA and incubated in the cells for 1 h at RT. The following primary antibodies were used: vimentin mouse antibody (dilution 1:150, Sigma, V6630); and vinculin mouse antibody (dilution 1:500, Sigma, V9131). Secondary antibodies were used: Goat anti-Mouse IgG (H+L) Fluor™ 555 (dilution 1:1000, Invitrogen, A-21422), wheat germ agglutinin (WGA, dilution 1:700, Invitrogen, W11261), DAPI Fluoromount-G reagent (SountherBiotech, 0100-20) was used to mount cells on cover slide and imaged using Olympus spinSR10 Ixplore spinning disk confocal microscope.

### Nuclear-cytoplasmic fractionation

MCF-7 cells treated with THP-1 supernatant were collected and resuspended in PBS, distributed evenly into two tubes. After centrifugation at 800 rpm for 5 min, the supernatant was discarded. For fractionation, cells were lysed on ice in 120 µL of HEPES buffer (10 mM HEPES, 10 mM KCl, 1.5 mM MgCl_2_, 0.2% NP-40, pH 7.5) for 30 min, followed by centrifugation at 5000×g for 10 min at 4°C. The supernatant was collected as the cytoplasmic fraction and mixed with 30 µL of 5× loading buffer. The pellet was washed three times with 200 µL of HEPES buffer and then lysed in RIPA buffer on ice. After centrifugation at 13,400×g for 10 min at 4°C, the supernatant was mixed with 30 µL of 5× loading buffer to obtain the nuclear lysate.

### Real-time RT-PCR

Total cellular RNA was extracted by EZ-press RNA Purification Kit (EZBioscience, B0004DP) according to the manufacturer’s protocols. Total RNA was reverse transcribed by using ColorReverse Transcription Kit (EZBioscience, A0010CGQ). Real-time RT-PCR was carried out by using 2× Color SYBR Green qPCR Master Mix (ROX2 plus) (EZBioscience, A0012-R2) in QuantStudio 1 system (Thermo Scientific). All data were normalized to the GAPDH expression level. Primer sequences used for real-time RT-PCR were as follows: *ITGB6*: Forward 5’-TCTGGAGTTGGCGAAAGG-3’, Reverse 5’- TCCACCGGGTAGTCCTCA-3’, *ITGAV*: Forward 5’- ATCTGTGAGGTCGAAACAGGA-3’, Reverse 5’- TGGAGCATACTCAACAGTCTTTG-3’, *VIM*: Forward 5’-GACCTTGAACGCAAAGTGGAATC-3’, Reverse 5’-GTGAGGTCAGGCTTGGAAACATC-3’, *GAPDH*: Forward 5’-GCATCCTGCACCACCAACTG-3’, Reverse 5’- GCCTGCTTCACCACCTTCTT-3’.

### Drugs application

Inhibitors used in this study included Z-VAD-FMK (pan-caspase inhibitor, MedChemExpress, HY-16658B) at 30 μM for 24 h; Z-DEVD-FMK (caspase-3 inhibitor, APExBIO Technology LLC, A1920) at 20 μM for 24 h; Monensin (classical secretory pathway inhibitor, Beyotime, S1753) at 3 μM for 4 h; GW4869 (exosome inhibitor, MedChemExpress, HY-19363) at 10 μM for 24 h; Punicalagin (non-classical type I secretion pathway inhibitor, MedChemExpress, HY-N0063) at 5 μM for 1 h; glycine (plasma membrane rupture inhibitor, Sigma, G7126) at 50 μM for 10 min; GRGDSP (pan-integrin inhibitor, MACKLIN, G876434) at 250 μM for 1 h; EMD527040 (Integrin αVβ6 inhibitor, MedChemExpress, HY-101473) at 3 μM for 24 h; Picropodophyllin (PPP, IGF-1R inhibitor, MedChemExpress, HY-15494) at 1 μM for 24 h; FMK (RSK inhibitor, APE BIO, A3420) at 3 μM for 3 h; IGF1 (IGF-1R ligand, Novoprotein, C032) at 50 μM for 20 min. Peptide 71-90, corresponding to amino acids 71-90 of the VIM protein, was synthesized (Sangong) as follows: RSSVPGVRLLQDSVDFSLAD, at 10, 50, and 100 μM for 24 h. Purified FL VIM was used at 20 μM for 24 h.

### Human phospho-kinase array

5×10^6^ cells were seeded in a 24-well plate and allowed to adhere for 24 h, and THP-1 supernatant or vimentin KO THP-1 was added. The experiment was performed according to the manufacturer’s instructions provided with the kit (R&D Systems, ARY003C).

### Cell viability assay

Cell growth curve and viability were measured by CCK-8 kit (Beyotime, C0039), and lactate dehydrogenase (LDH) release was measured by LDH kit (Beyotime, C0016), following the manufacturer’s protocol.

### Immune cell infiltration analysis

The RNA-sequencing expression profiles and associated clinical data for breast cancer (BRCA) patients at late stages (vi-x) were obtained from the Cancer Genome Atlas (TCGA) dataset (https://portal.gdc.com). The immune score assessment, indicative of the level of immune cell infiltration, was conducted using R software packages including devtools, e1071, preprocessCore, parallel, bsqsc, ggplot2 and CIBERSORT. The aforementioned analytical approaches and R package were executed using the R foundation for statistical computing (2020) version 4.0.3.

## Consent for publication

All authors consent to the publication of this study.

## Availability of data and material

All data and material generated in this study will be freely provided upon request to the corresponding author.

## Acknowledgments

This work was supported by National Natural Science Foundation of China (92354301, 32222022, 92054104); R&D Program of Guangzhou National Laboratory (GZNL2023A03004); Natural Science Foundation of Shanghai (23ZR1470900); Key Research and Development Program, Ministry of Science and Technology of China (2022YFC2303502, 2021YFC2300204).

## Author contributions

C.M. and X.G. carried out the majority of the experiments and interpretation of the data. J.M., C.W. and Y.H. carried out the protein purification experiments. W.W. and S.Z. performed the AlphaFold prediction. Z.L. and X.H. participated in the *in vivo* experiments. Z.C. participated in preparing the human patient samples. Z.W. and X.L. participated in the RNA sequencing and analysis. G.F., C.T., X.L. and C.W. participated in experiment design and supervision. Y.J. and T.X. conceived and supervised the project. Y.J., C.M. and X.G. wrote the manuscript with contributions from all other authors.

