## Supplemental figures for "A novel vimentin variant from tumor-associated macrophages directs cancer metastasis by engaging IGF-1R"

#### Extended Data Fig. 1

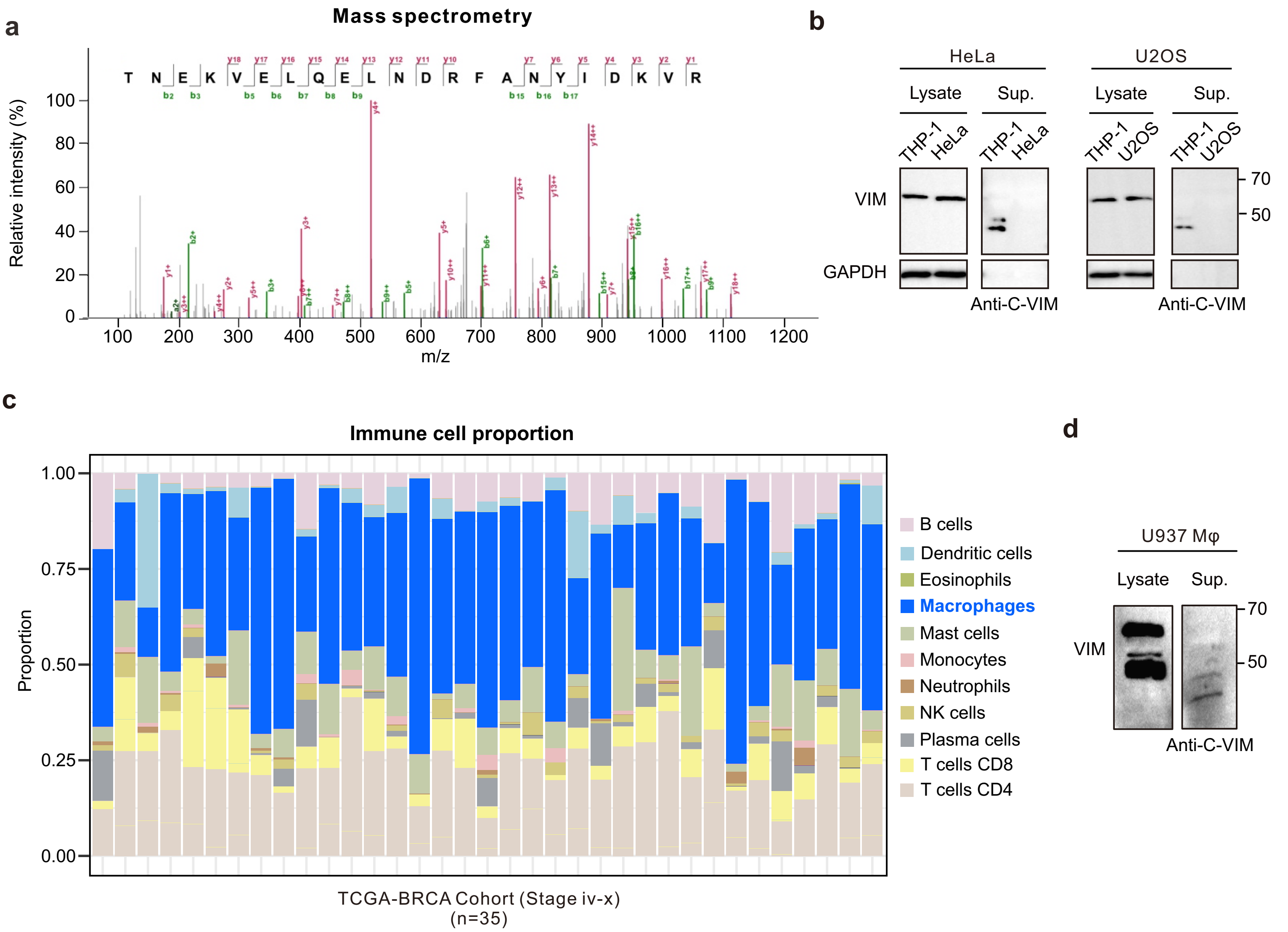

Extended Data Fig. 2

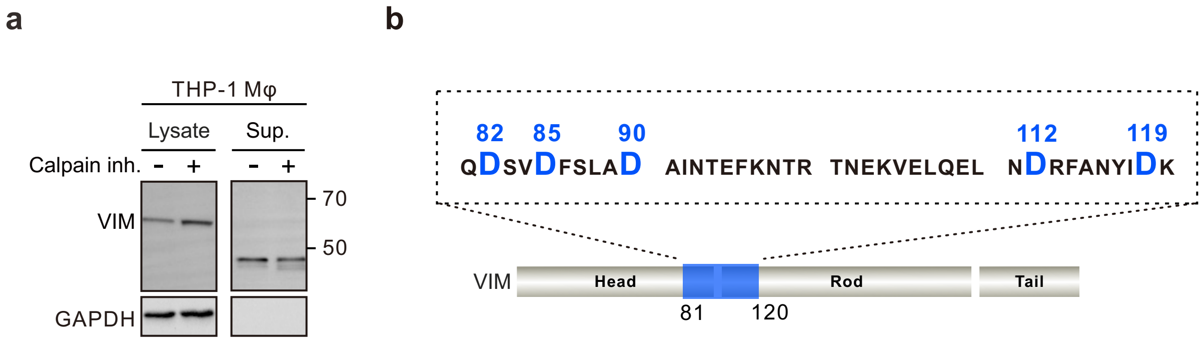

#### Extended Data Fig. 3

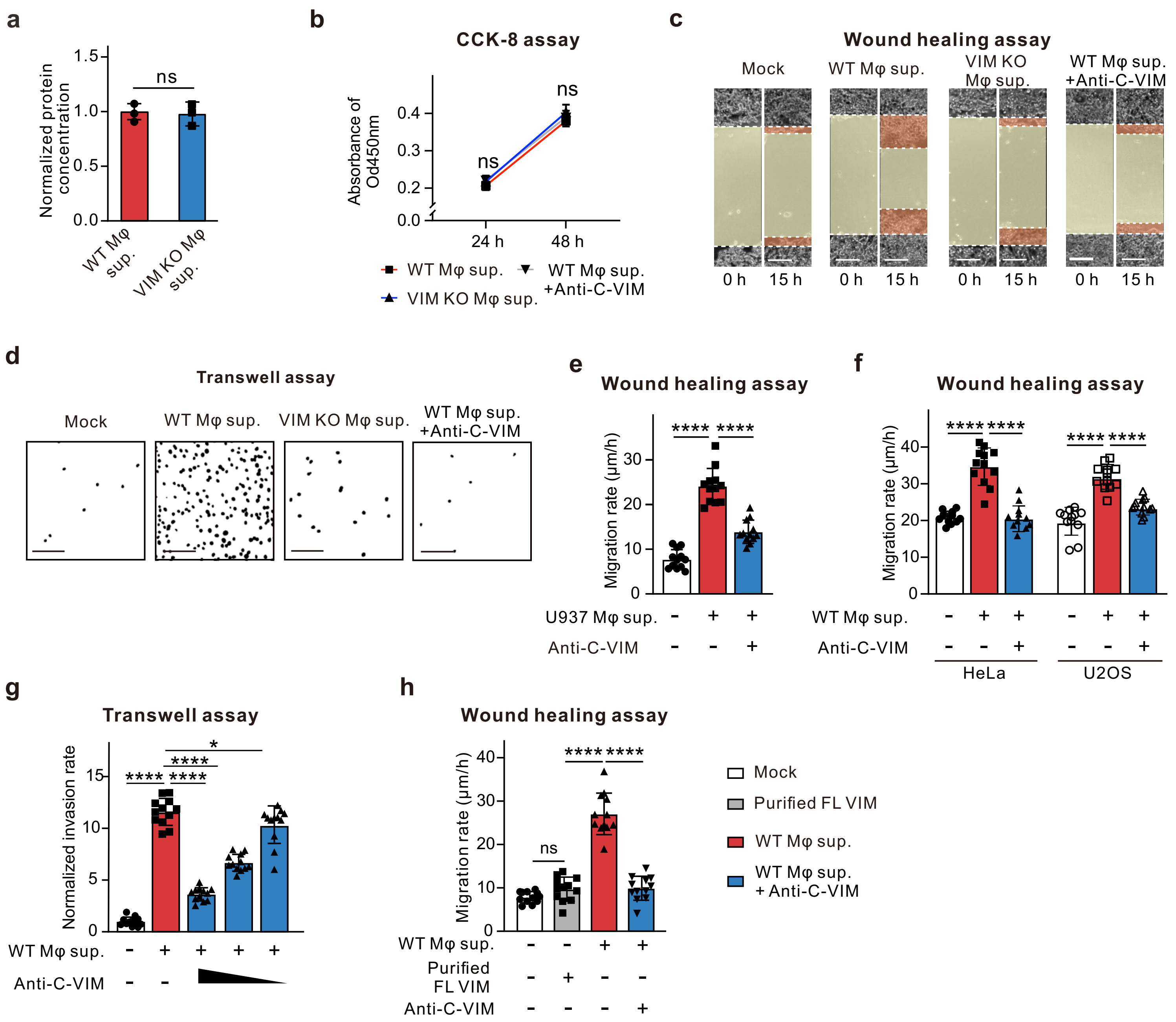

#### Extended Data Fig. 4

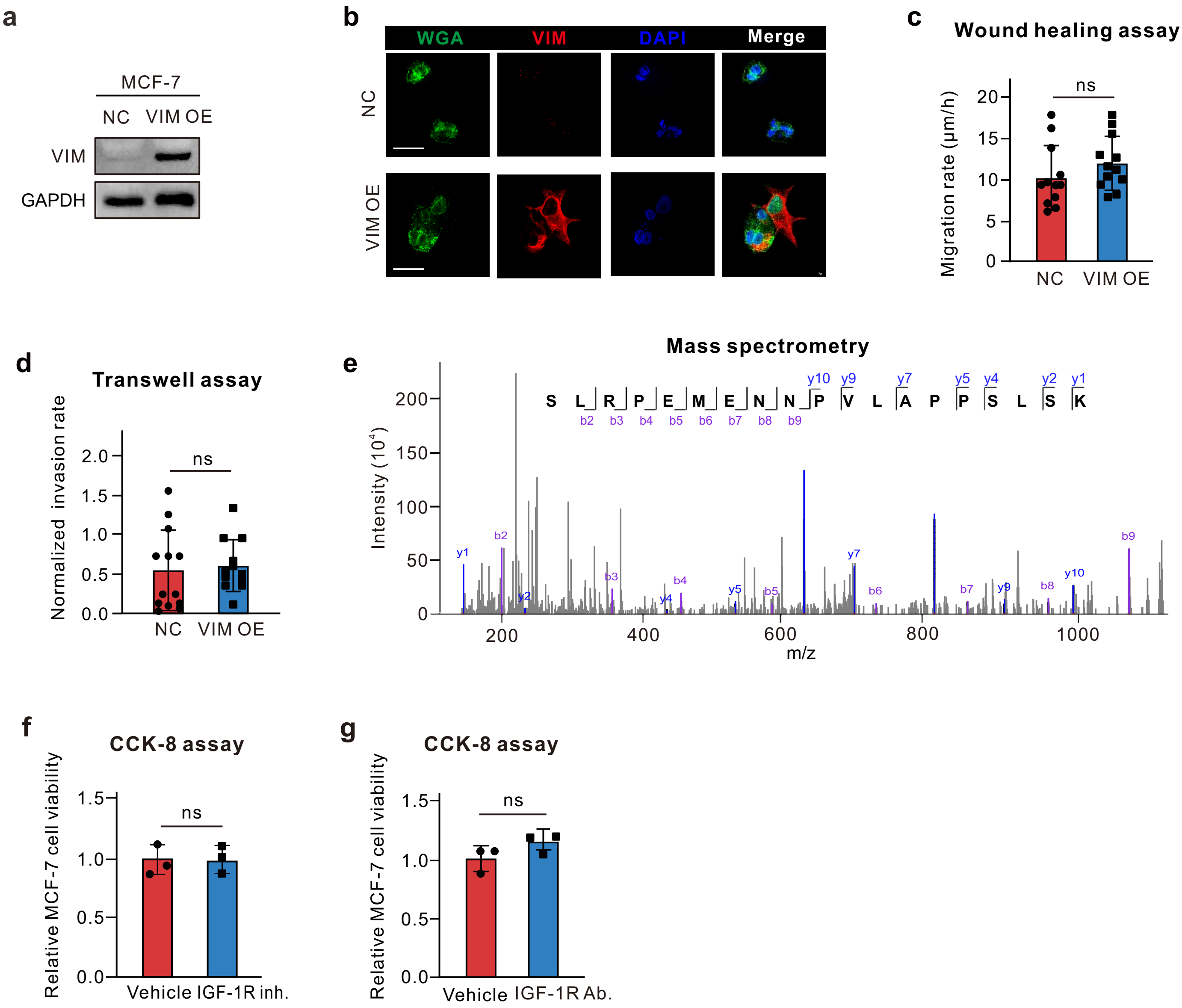

#### Extended Data Fig. 5

**a**

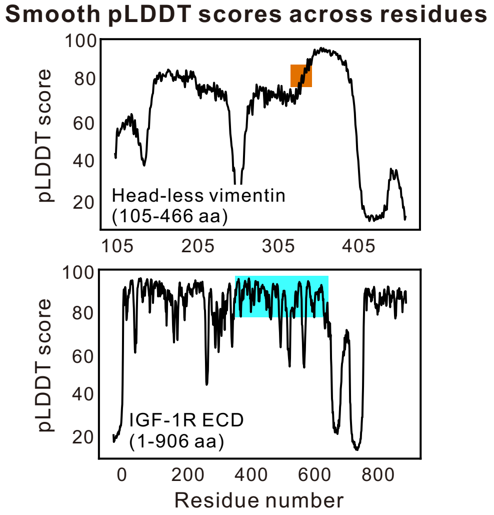


**b**

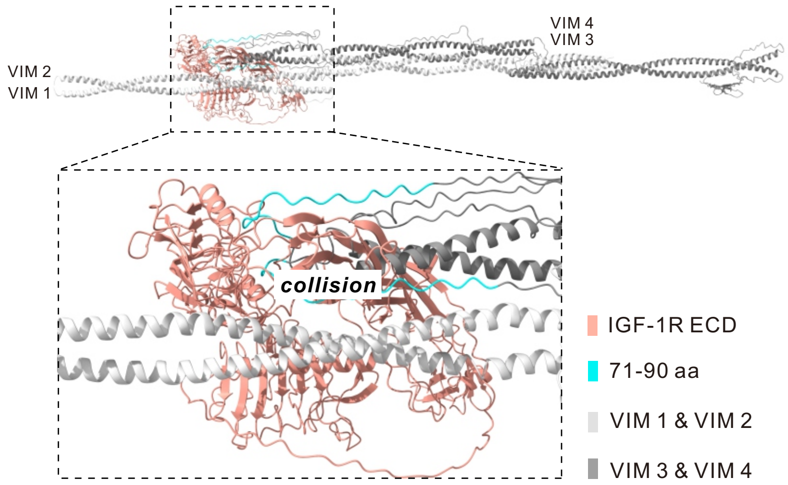


**c**

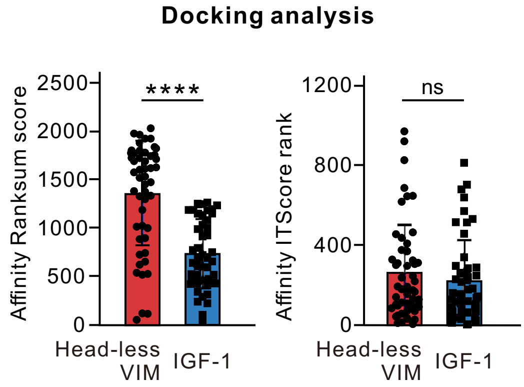


**d**

| | mssVIM | IGF-1 |
|---|---|---|
| Source | TAM exclusive | Various |
| Biological effects | Cancer cell migration | Proliferation Inflammation |
| Affinity to IGF-1R | Ratively strong | Strong |

#### Extended Data Fig. 6

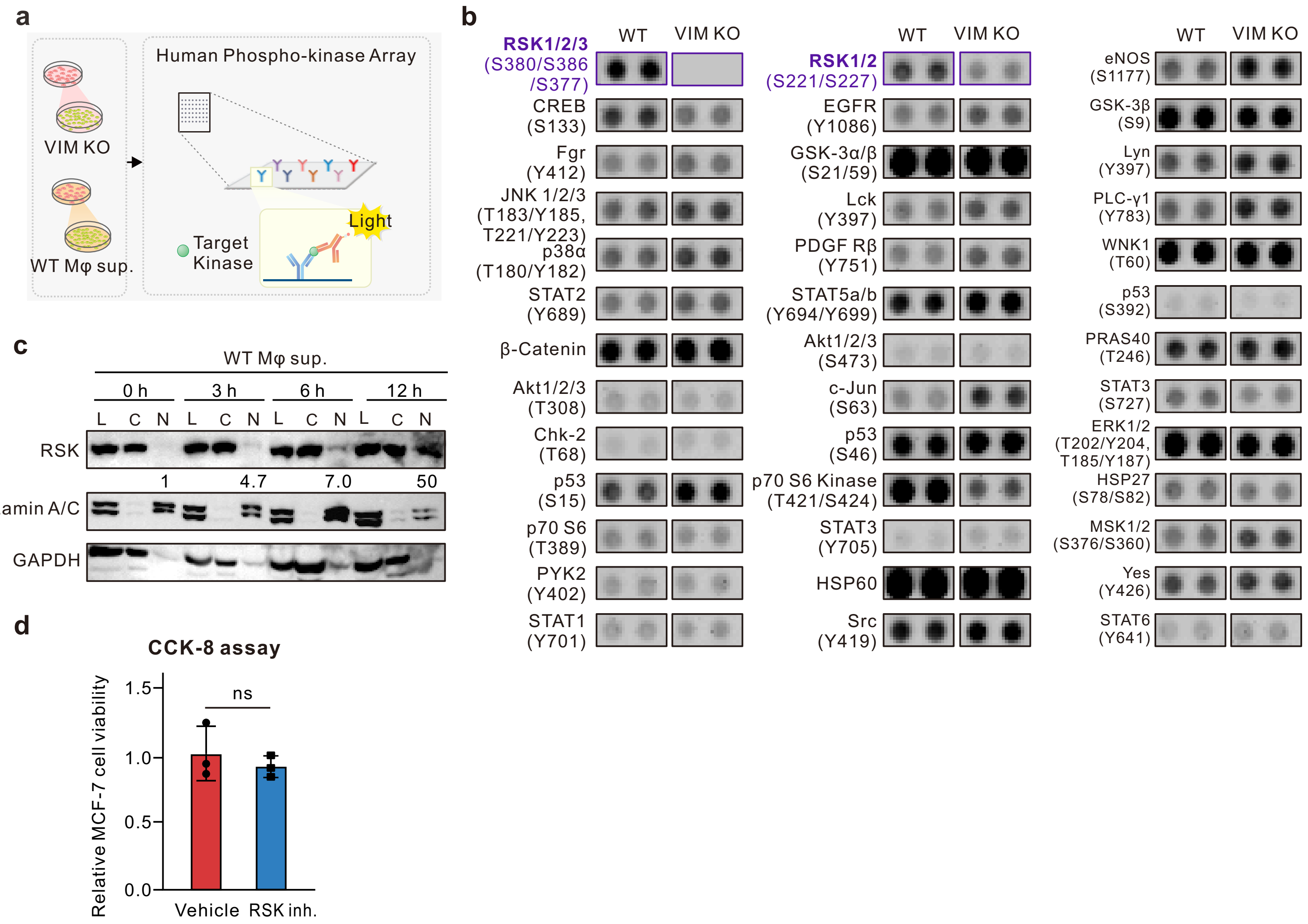

### Extended Data Fig. 7

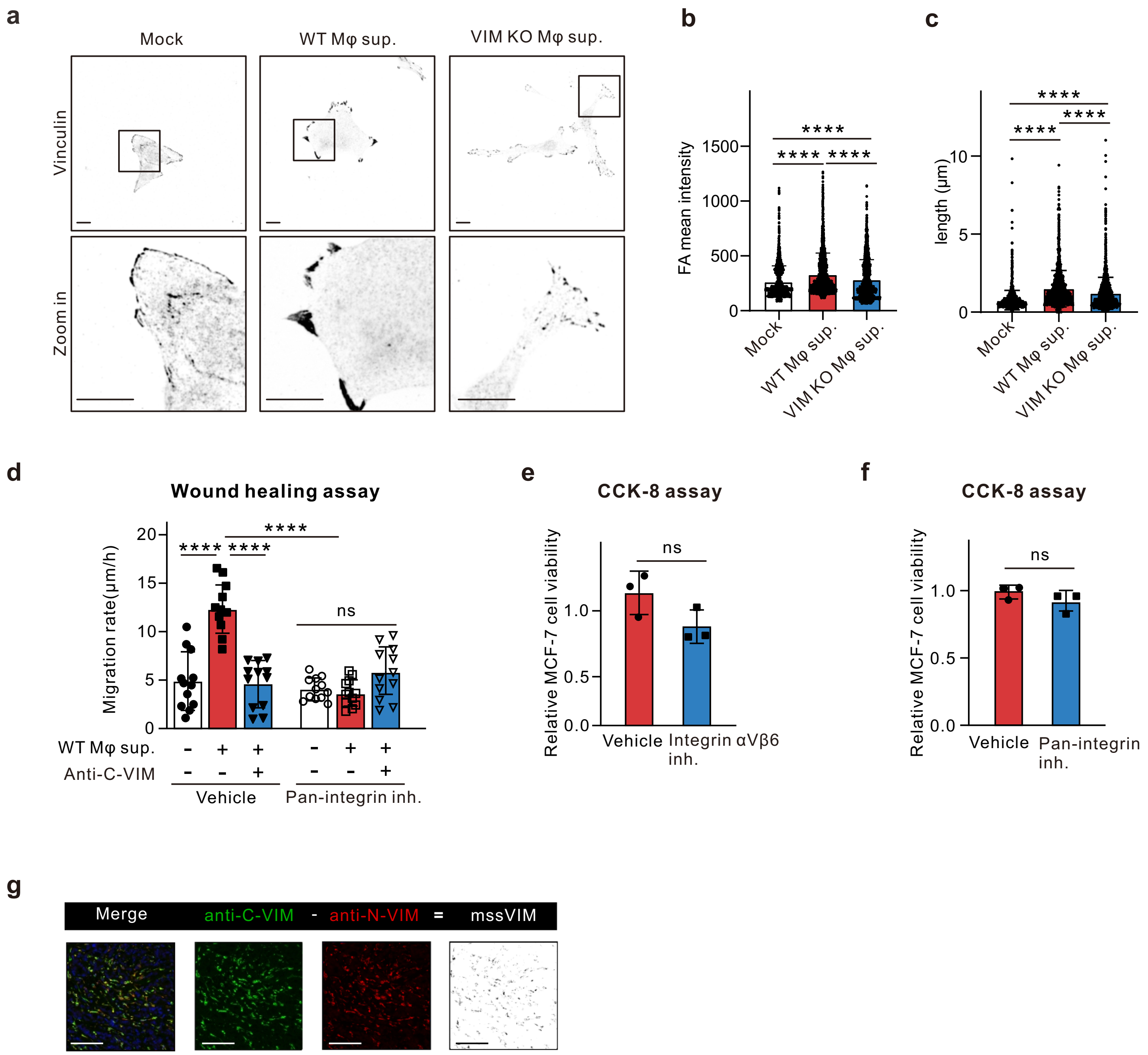
